# Artificial selection for resistance to copper and off-target physiological and behavioral effects in *Drosophila melanogaster*

**DOI:** 10.1101/2025.09.30.679338

**Authors:** Kazzrie Arnold, Elizabeth R. Everman

## Abstract

Pollution resulting from mining, industry, and agriculture has played an important role in shaping the evolution of diverse organisms. The pathway to evolved resistance can involve polygenic shifts that affect multiple traits. Heavy metal contamination is of particular concern due to the health effects for humans and cross-tolerance effects that influence pesticide resistance. We used a replicated artificial selection approach to examine the response to copper selection in *Drosophila melanogaster* collected from a retired mine and active fruit orchard. We tracked shifts in resistance to the target trait, copper, as well as off-target effects on cadmium and lead resistance, starvation resistance, lifespan, and feeding aversion to copper contaminated food. Selection for copper resistance increased the focal trait and maintained resistance to non-essential heavy metals while control flies lost metal resistance overtime, implying a cost associated with maintaining resistance to heavy metals. Starvation resistance was correlated with copper resistance, but the correlation was not sufficient to explain the gain in copper resistance. Lifespan was correlated with increased copper resistance in flies collected from one of the two collection sites, suggesting that life history traits may be influenced by repeated heavy metal exposure. Future genomic analysis will help clarify the genetic control of the selection response. Together, our results underscore the complexity of adaptive shift in polygenic traits and provide a basis for further exploration of costs and correlative change following copper selection.

## Background

Resistance to chemical stressors has repeatedly evolved in response to mining, agriculture, industry, and other anthropogenic sources in a wide range of organisms (1–8). In some cases, the evolved response is accompanied by cross-tolerance (a correlated increase in resistance) to other environmental stressors (9–13) and in other cases changes in life history traits that affect fitness (14–16). For sources of stress that are entirely xenobiotic (not required for normal physiological function, development, or metabolism), adaptive response has been repeatedly shown to influence single or few major effect loci (8,17–19) (although this pattern is not canonical, for example see (20)). This simpler pattern of adaptive response is sometimes framed in contrast to a more polygenic pattern of adaptation of complex traits, where polygenic traits shift over time due to changes at many smaller effect loci that are distributed across the genome, in effect offering many unique or shared paths to the optimal trait (17,21–27).

Resistance to potentially toxic substances that are biologically necessary at low levels offers an opportunity to explore the process of complex trait adaptation and its potential consequences for other traits. The heavy metal copper is a conducive model stressor for this frame: copper is an essential micronutrient that plays a critical role in cellular respiration and the oxidative stress response (28–31), copper is toxic at high levels and can result in tissue and neurological damage (29,32), and resistance to copper toxicity is genetically complex (33), varies in natural populations, and is responsive to environmental exposure (1,2,7,34,35). That there are multiple adaptive routes to copper resistance has also been empirically demonstrated. Experimental evolution of copper resistance using 34 *Saccharomyces cerevisiae* strains identified over 50 unique genetic changes (SNPs, CNVs, and aneuploidy) associated with increased copper resistance, ultimately demonstrating that multiple combinations of genetic changes can confer resistance (36). Given the high number of potential loci that can contribute to copper adaptation and the overlap in the genes responsible for regulating, metabolizing, and detoxifying heavy metals that are necessary (e.g. Cu, Zn) or always toxic (Cd, Pb) (37,38), copper selection has the potential to result in cross-tolerance to other stressors or result in correlated shifts in fitness-relevant traits.

In the present study, we examined the target and off-target effects of artificial selection for resistance to copper sulfate in *Drosophila melanogaster*. *D. melanogaster* is well suited for this endeavor because of its short generation time and ease of copper exposure through food. We have also previously characterized the genetic architecture of survival following copper exposure using two large laboratory mapping populations (the *Drosophila* Synthetic Population Resource, DSPR (39) and the *Drosophila melanogaster* Genetic Reference Panel, DGRP (40)) as well as multiple populations sampled from agricultural and mining sites (33,35,38,41). In our previous work that examined variation in copper resistance among wild *D. melanogaster* populations, we identified a high copper resistance population inhabiting a retired copper mine (Burra Burra Mine, BBM) in the Copper Basin of Tennessee, USA where mining occurred for six decades. Bulked-segregant analysis of flies from the BBM site revealed several candidate loci that have been previously linked to mitochondrial function, oxidative stress, arsenic and lead response, and ATP synthesis (35). The broad nature of the candidate gene categories raises the question of whether copper resistance can be further increased in the BBM population compared to another non-mine population, whether copper resistance correlates with resistance to other stressors, and whether there are implications for behavioral and life history traits as a result of evolved copper resistance.

In general, heavy metal toxicity results in oxidative stress and the displacement of biologically necessary metal cofactors with toxic metal ions (42–44). Copper, zinc, lead, cadmium, and mercury have all been shown to illicit these effects (45,46), and genes such as the Metallothioneins are upregulated in response to several heavy metals (38,46–49). We expect that increased resistance to copper toxicity will be correlated with increased resistance to non-essential heavy metals such as lead and cadmium. In addition, given that heavy metal toxicity can have broad physiological effects (i.e. oxidative stress, activation of heat shock proteins (50)) and previous associations between copper and starvation resistance (35), we expect copper selection will also increase starvation resistance. Further, the genetic shifts that result in increased copper resistance may have effects on fitness traits related to life history (51) For example, selection for cadmium resistance in *Nilaparvata lugens* increased insecticide resistance but had a varied fitness effects, reducing fecundity, delaying development, but increasing lifespan of females (13). Finally, organisms interact with their environment via sophisticated chemosensory organs that can influence the consumption of contaminated food. For example, evolved responses to insecticides can be considered a composite of physiological and behavioral responses (52,53). However, outside of the effects of heavy metal stress on memory and cognition (54,55), the impact of selection for resistance to heavy metals on behavior has not been investigated.

Using flies collected from BBM and a nearby agricultural site in the Copper Basin region (Mercier Orchards (MO), Blue Ridge, GA, USA), we examined the effect of artificial selection for copper resistance on the target trait (adult copper resistance) as well as cadmium, lead, and starvation resistance using a replicated experimental evolution design (Figure S1). We also tested the effect of copper selection on the behavioral trait feeding preference to determine whether aversion to copper in food responded to selection. To investigate the potential effects of copper selection on life history, we measured variation in lifespan. Overall, we found a clear positive effect of selection on adult copper resistance. Resistance to non-essential heavy metals lead and cadmium did not increase but was maintained in flies from selected cages. Control cages that were not subjected to selection lost cadmium and lead resistance over time. Consistent with patterns observed in wild populations (35), starvation resistance was significantly correlated with copper resistance, but resistance to copper was not fully explained by increased aversion to copper-containing food. Instead, all flies (control and selected) gradually lost their aversion to copper over time. Finally, we observed an association between adult copper resistance and lifespan that varied depending on the source of the wild-derived flies. Flies from the BBM site retained longer lifespan regardless of selection, while MO-derived flies gained longevity in association with copper resistance. Future work will examine the genetic shifts through artificial selection to characterize overlap between traits.

## Materials and Methods

### Collections and rearing

Between July 09 and July 18, 2024, we collected gravid *Drosophila melanogaster* females daily from two locations (Burra Burra Mine, BBM and Mercier Orchards, MO; Table 1), which reside in or near the Copper Basin region of the United States.

**Table 1.** Collection sites and number of females collected from each site.

| Site | Location | Collection Location | N Founder Females |
| --- | --- | --- | --- |
| Burra Burra Mine (BBM) | 35.04 N, -84.38 W | Apple tree | 191 |
| Mercier Orchards (MO) | 34.89N, -84.34W | Dumpsters, cider vats, blackberries, unidentified deciduous trees | 355 |

To collect flies, traps made from plastic bottles (ULINE, S-21727W) were prepared and baited with banana and yeast as described in Everman et al. (35). Traps were suspended from structures (tree branches, railings) and were allowed to ferment for up to three days before replacement. In addition to passive trap collection, we used sweep netting to collect flies hovering near fruit sorting bins at the MO site and near dropped apples at the BBM site. We mouth-aspirated single female flies from traps or nets into cornmeal-molasses-yeast vials to collect eggs for species verification as described in Everman et al. (35).

Generation 0 (G0) offspring of the species-verified *D. melanogaster* females were used to establish generation 1, from which five G1 males and females per original founder female were used to establish one population cage (30×30×30cm, Bugdorm-1) each for BBM- and MO-derived flies. Eggs were collected from G2 individuals in cages following Everman et al. (35) and were pipetted into 24 6oz *Drosophila* bottles (Genessee) containing media following the Bloomington *Drosophila* Stock Center Semi-Defined Food formulation (56). Bottles containing approximately 500 eggs each were used to establish six cages per collection site, resulting in a total of 12 wild-derived G3 cages (Figure S1). All cages were maintained in the same Percival incubator at 25°C, 50% relative humidity, and under a 12:12 L:D cycle throughout the artificial selection process and for all phenotyping.

### Baseline phenotyping of wild-derived flies

In addition to establishing the cages for MO and BBM-collected flies, we also maintained the offspring of each founder female as isofemale strains in vials. Using G2 individuals, we measured Adult Copper Resistance (ACR) and Adult Starvation Resistance (ASR) to estimate the baseline levels of resistance in the MO and BBM collection sites in flies as close to field-caught as possible and prior to artificial selection (Figure S1). Flies were sorted into groups of 20 individuals over CO_2_ and allowed to recover for 24 hours on control food. Flies (3-5 days old) were then transferred to vials containing 1.8g Instant *Drosophila* Media (Carolina Biological Supply Company 173200) hydrated with 50mM CuSO_4_ (Copper(II) Sulfate; Sigma-Aldrich C1297) or starvation media (1.5% agar with media preservatives as described in Everman et al. (57). Survival was assessed every 24 hours until all flies were dead. ACR and ASR were measured in 20 flies/sex/founder female/collection site.

### Artificial selection and phenotyping of wild-derived population cages

Following the establishment of the replicated G3 population cages, we collected G4 eggs from each of the 12 cages (Figure S1) into vials to obtain estimates for each of the traits that would be tracked through artificial selection. Unless otherwise noted, eggs were dispensed in aliquots of 30uL onto cornmeal-molasses-yeast media. Eggs were also collected into bottles (semi-defined media) to establish the next generation of cages (Figure S1).

Selection for adult copper resistance occurred in every other generation, starting with G5 adults. Following emergence of adult flies in each cage, source bottles were removed and replaced with bottles containing semi-defined media with 50mM CuSO_4_ in each selection cage (4 bottles/cage). Control cages received four bottles of uncontaminated semi-defined media to prevent starvation. Once approximately 50% of flies had died in the selection cages, we added another set of control bottles to all cages, including control cages to prevent starvation (8 bottles/selection cage, 4 bottles/control cage). We found that females typically laid fewer eggs after copper exposure, making apple juice plate egg collection in selection cages inefficient. Instead, we allowed females to lay eggs directly into the eight bottles to ensure a large enough population size in the next generation. In contrast, control cage females laid large numbers of eggs very quickly, necessitating collection of eggs with apple juice plates to control density.

Once egg collection was complete, adults from the selection generation were frozen and discarded to ensure non-overlapping generations. Cages were cleaned with warm water to remove all flies and bottles with even-numbered generation flies were placed into their respective cages. Once even-numbered generation flies emerged, we repeated the entire process through Generation 20 (8 selection events) as follows: We collected eggs into vials for phenotyping and collected eggs into bottles for the next generation, which would be exposed to adult selection. We collected eggs from the generation following selection to decouple adult exposure from developmental effects on the offspring used for trait estimation (described below).

### Adult stress response

Three days following first emergence from vials, male and female flies were separated and sorted into groups of 20 individuals over light CO_2_ anesthesia and allowed to recover for 24 hours. Following recovery, 3-5 day old flies were transferred to experimental vials containing 50mM CuSO_4_ (**ACR**; 5 vials/sex/cage), starvation media (**ASR**; 5 vials/sex/cage), 20mM CdCl_2_ (Cadmium Chloride Sigma 655198; **ADR**; 3 vials/sex/cage), or 100mM Pb(CH_3_CO_2_)_2_ (Lead(II) Acetate Trihydrate, Sigma 228621; **ALR**; 3 vials/sex/cage). The lead and cadmium food were prepared in smaller quantities and with lower replication to reduce the impact of our experiments. We used 1/8^th^ of a teaspoon of finely ground Instant *Drosophila* media and hydrated food with 900uL of either metal in small plastic caps (MOCAP FCS131/16NA1) that fit within the narrow *Drosophila* vial opening. Caps were held onto vials with tape and disposed up following University guidelines. ACR and ASR vials were prepared as described above. Dead flies in each vial were counted daily to estimate survival and average lifespan (hours) per vial, per cage.

### Behavioral response to copper

We measured adult preference/avoidance of copper contaminated food (CA) using the Microplate Feeding Assay developed by Walters et al. (58) in both males and females (3-5 days old, 36 flies/sex/cage). This assay involved a 96-well plate containing 100uL of starvation media and a 3D printed coupler, which formed a connection between the starvation plate and a 1536-well plate that contained 10uL liquid food dyed with 40ug/mL erioglaucine disodium salt (Sigma 861146) and different concentrations of copper. Flies were placed into the starvation plate (1 fly/well of rows 2-7 of each plate) and allowed to recover from CO_2_ and to fast for 24 hours. Liquid food was prepared according to Walters et al. (58) except that instead of ethanol, we tested 0 (control), 0.5, 1, and 2mM CuSO_4_. The quantity of food consumed was estimated by comparing absorbance of food prior and following the feeding assay. We obtained initial absorbance readings at 630nm for each 1536-well plate, allowed flies access to the food for 24 hours, and then remeasured absorbance for each plate. After correcting for evaporation (58) and removing observations for flies that had died or not eaten food during the assay, we calculated preference per fly as (uL Copper Consumed – uL Control Consumed)/(Total uL Consumed). Because we calculated preference relative to the copper consumption, negative values indicate copper avoidance, and positive values indicate copper preference.

### Adult lifespan

Flies were collected for lifespan estimates (ALS) at the same time and in the same manner as for adult stress response traits. We measured adult lifespan in G4, G8, G12, G16, and G20 (3 day old starting age, 10 flies/5 vials/sex/cage). Flies were held on cornmeal-molasses-yeast media throughout the duration of their lifespan and were transferred to new vials on Monday, Wednesday, and Friday of each week. Counts of survival were made on each day that flies were transferred to new food. Average lifespan (days) was estimated for each vial per cage.

### Data analysis

All data preparation and analyses were carried out in R (V 4.5.1) (59) and RStudio (V 2025.5.1.513) (60). Significance was assessed at an alpha level of 0.05, and experiment-wide alpha levels were controlled for post hoc tests. Data visualization was accomplished using ggplot2 (61).

#### Baseline G2 assessment of ACR and ASR

After calculating average lifespan on either treatment per vial of 20 flies, data were analyzed with a two-way ANOVA (Average Lifespan ∼ Sex * Collection Site) for each trait. Because we measured only one vial of females and males per founder female strain, the strains were treated as replicate estimates of each of collection site. We also assessed the correlation between ACR and ASR, accounting for the effects of sex and collection site with multiple regression (ACR ∼ Sex * Collection Site * ASR) using the lm function.

#### G2 versus G4 assessment of ACR and ASR

The transition of flies collected from the field to vials and cages maintained in the lab has the potential to influence phenotypic variation through bottlenecks and through rapid lab adaptation. To assess the starting levels of ACR and ASR in G4 caged flies prior to artificial selection for copper resistance relative to the G2 vial-level assessment, we used a three-way ANOVA to test the effects of container (cage vs. vial), sex, and collection site (ACR or ASR ∼ Sex * Collection Site * Container) to determine if ACR and ASR were different in flies maintained in cages versus vials.

#### Effect of selection and trait correlations

To assess the influence of selection on the change in each trait, we used analyses of covariance (ANCOVAs), accounting for cage, collection site, selection regime, treatment (when applicable), sex, and their interactions. Generation was treated as a covariate in each analysis and interactions between model terms and generation were used to determine whether the slopes of each response were different. Post hoc tests of significant terms were completed using estimated marginal means (emmeans_test) with Bonferroni correction using the R package rstatix (62). Following initial analysis of ACR data, we found systematic bias in G6 data that was attributed to differences in experimenter handling of flies, so this generation was removed from analysis.

Correlations between selection cage-level estimates of copper resistance and all other traits were also carried out with ANCOVAs as well, accounting for model terms as described above and excluding G6. We also estimated Pearson Correlations using the rcorr function from the R package Hmisc (63).

#### Assessment of effect of selection versus G2 flies

We compared the adaptive response of MO and BBM flies to their respective G2 estimates of ACR using a three-way ANOVA, testing the effects of sex, generation (G2 vs G20), and collection site using data from the selection cages. Differences among G20 estimates of ACR between selection cages were determined with a three-way ANOVA (testing effects of sex, collection site, and cage) for G20 data followed by Tukey HSD Post hoc comparisons.

### Data availability

All phenotype data generated in this study are available from FigShare (10.6084/m9.figshare.29971417).

## Results

### Baseline ACR and ASR differed between collection sites

Our two collection sites are environmentally distinct and have experienced different types of anthropogenic impact (retired mine vs. active agriculture). To characterize baseline Adult Copper Resistance (ACR) and Adult Starvation Resistance (ASR), we measured both traits in offspring descended from founder females collected from each site.

Baseline ACR was highly variable in both collection sites, ranging from 51.6-120hrs in BBM females (48-88.8hrs in BBM males) and from 50.5-124.0hrs in MO females (44.6-86.4hrs in MO males). Flies derived from the BBM collection site had slightly higher ACR compared to MO flies (Collection Site: F(_1,679_) = 7.65, P < 0.006; Figure 1A; Table S1). Males from both collection sites were consistently more sensitive to copper than females (Sex: F_(1,679)_ = 553.29, P < 0.00001; Collection Site x Sex interaction: F_(1,679)_ = 0.06, P = 0.8; Figure 1A). Overall, the similarity in ACR between BBM- and MO-derived flies suggests that both collection sites have experienced selection for copper resistance. If the minor difference in ACR between BBM and MO is considered biologically relevant, selection for copper resistance may be slightly higher at the BBM site.

**Figure 1.**
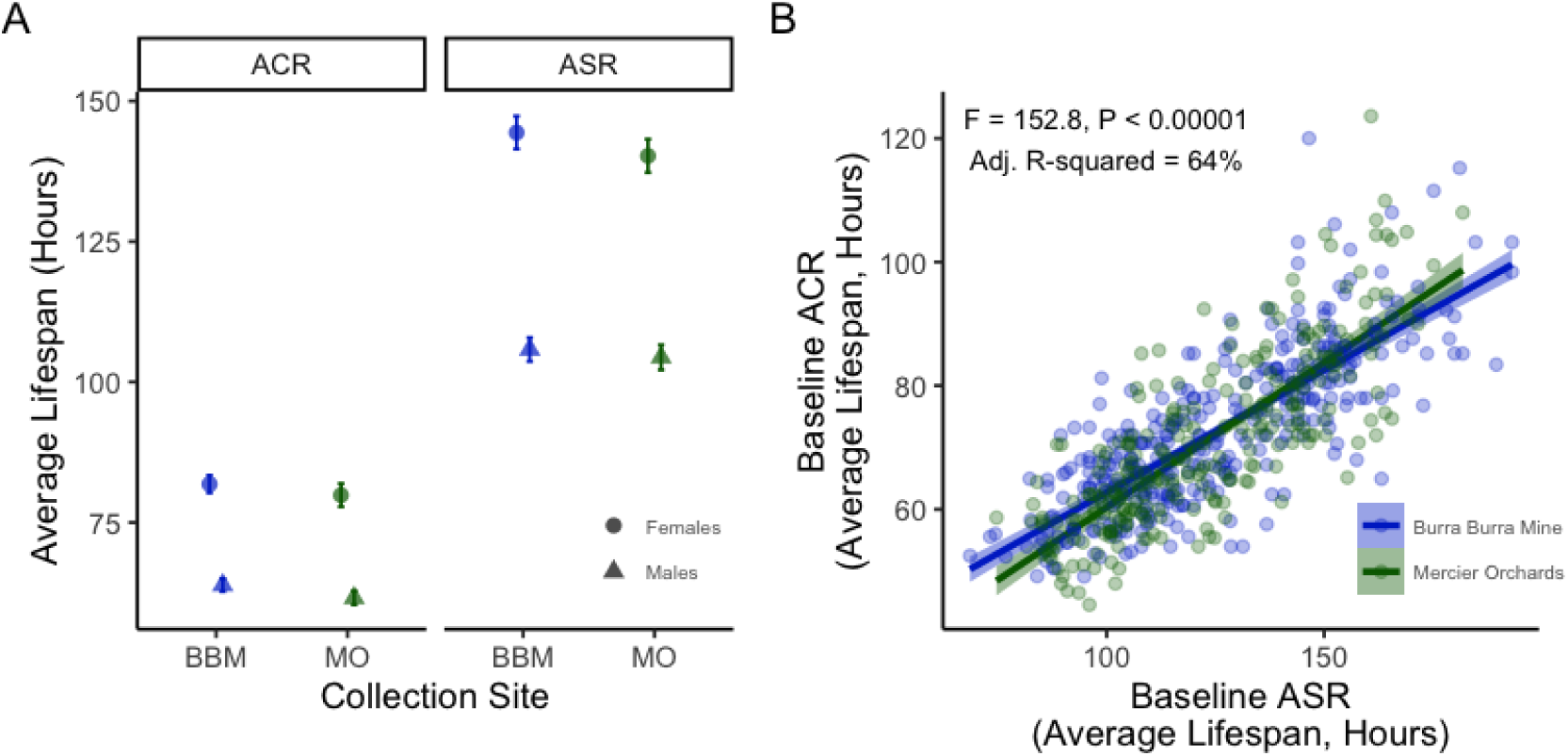
Baseline ACR and ASR in BBM and MO descended G2 flies. A. Baseline ACR varied due to population and sex, with flies from BBM capable of surviving on 50mM CuSO_4_ slightly longer than MO flies (P < 0.006). Females consistently survived longer than males (P < 0.00001; Collection Site x Sex interaction: P = 0.81). Baseline ASR also varied between BBM and MO (P < 0.04), and females were consistently more starvation resistant compared to males (P < 0.00001; Collection Site x Sex interaction: P = 0.31). In both panels, data shown are means +/-95% CI. B. ACR and ASR were significantly positively correlated (P < 0.00001, Adjusted R^2^ = 64%). The relationship between ACR and ASR was slightly stronger in MO compared to BBM (ASR x Collection Site: t = 2.49, P < 0.02). Points indicate sex-specific isofemale strain means and the shading shows the 95% CI around the regression (solid line). Statistic are presented in Table S1.

Because flies in our study are exposed to copper in their food and may survive copper in part by avoiding consumption of contaminated food, we measured Adult Starvation Resistance (ASR) and tested the relationship between ACR and ASR. ASR was similarly variable in each population (BBM females: 86.1-193.0hrs, BBM males: 68.6-141.0hrs; MO females: 97.2-182.0hrs, MO males: 74.7-140.0hrs) and BBM flies were slightly more starvation resistant compared to MO descended flies (F_(1,591)_ = 4.69, P < 0.04; Figure 1A; Table S1). Females consistently survived much longer under starvation conditions compared to males (Sex: F_(1,591)_ = 793.61, P < 0.00001; Collection Site x Sex interaction: F_(1,591)_ = 1.03, P = 0.31; Figure 1A).

The relationship between baseline ACR and ASR, accounting for sex and collection site, was significantly positively correlated (F_(7,587)_ = 153, P < 0.00001, Adjusted R^2^ = 64%; Figure 1B). We also detected a small but significant interaction due to collection site, due to a slightly stronger correlation between traits in MO flies (ASR x Collection Site: t = 2.49, P < 0.02; Figure 1B). Because starvation resistance has the potential to evolve in response to selection for copper resistance, we continued to track this trait throughout the artificial selection experiment.

### Initial stress response was similar between replicate cages

The offspring of collected females progressed through four generations before they were maintained in the cages for selection. Despite efforts to prevent artifacts due to lab adaptation and the establishment of cages, it is possible that our replicate cages may have begun the artificial selection experiment with different starting levels of resistance. We used a three-way ANOVA to determine whether there were significant differences between each replicate cage and the G2 estimates obtained for ACR and ASR in MO- and BBM-derived flies.

All replicate populations were statistically indistinguishable from each other and from the G2 estimates of MO and BBM resistance (ACR, Container (Cage or Vial): F_(12,779)_ = 0.57, P = 0.87; ASR, Container: F_(12,691)_ = 0.31, P = 0.99; Figure 2; Table S2). Differences between BBM and MO collection sites in overall ACR resistance were still significant although ASR was not (ACR, Collection Site: F_(1,779)_ = 7.52, P < 0.007; ASR, Collection Site: F_(1,691)_ = 3.70, P = 0.05; Figure 2). The strong effect of Sex continued to be observed for each trait (ACR, Sex: F_(1,779)_ = 738.1, P < 0.00001; ASR, Sex: F_(1,691)_ = 1099.1, P < 0.00001; Figure 2). No interactions were significant (Table S2). Any lab adaptation that occurred in the first four generations of being maintained under laboratory conditions had no discernable effect on ACR.

**Figure 2.**
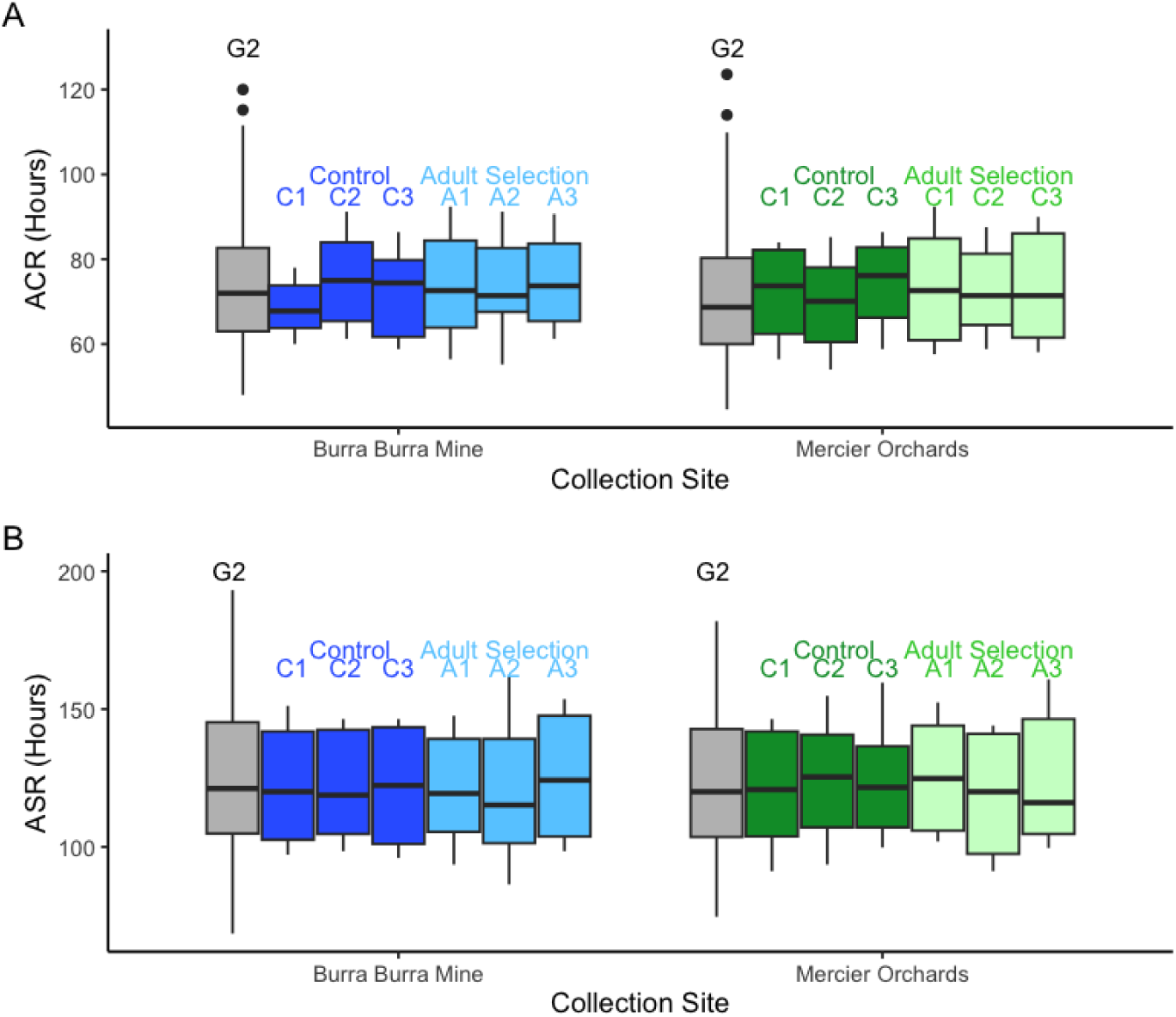
ACR and ASR in G4 flies maintained in cages prior to selection was not statistically different from G2 estimates. A. Copper resistance estimates in each of the replicate population cages planned for adult selection or control regimes were indistinguishable from each other and the G2 estimate (P = 0.87), although the ACR was slightly higher overall in all BBM-derived flies (P < 0.007). B. Starvation resistance was also indistinguishable among all collection site-specific cages (P = 0.99), and there were no collection site differences (P = 0.05). In both plots, BBM-derived estimates are shown in blue and MO-derived estimates are shown in green. Grey shading indicates G2 estimates; medium shading indicates the three replicate Control cages; lightest shading indicates the Adult Selection cages. Statistics are presented in Table S2.

### Selection increases adult copper resistance

Selection for copper resistance in the adult life stage began with adult G4 flies (separate from those used for phenotyping; Figure S1) and occurred in every other generation from that point on. Copper selection significantly increased ACR relative to control (non-selected) cages, with copper-selected flies surviving on average 16.7 hours longer than control flies on 50mM CuSO_4_ by G20 (Selection: F_(1,928)_ = 391.35, P < 0.00001; Selection x Generation: F_(1,928)_ = 98.69, P < 0.00001; Figure 3; Table S3). The overall effect of selection was similar between BBM- and MO-derived flies (Collection Site: F_(1,928)_ = 3.67, P = 0.06), and both BBM and MO cages responded to selection over time in a similar manner (Collection Site x Selection x Generation: F_(1,928)_ = 0.0010, P = 0.97; Figure 3).

**Figure 3.**
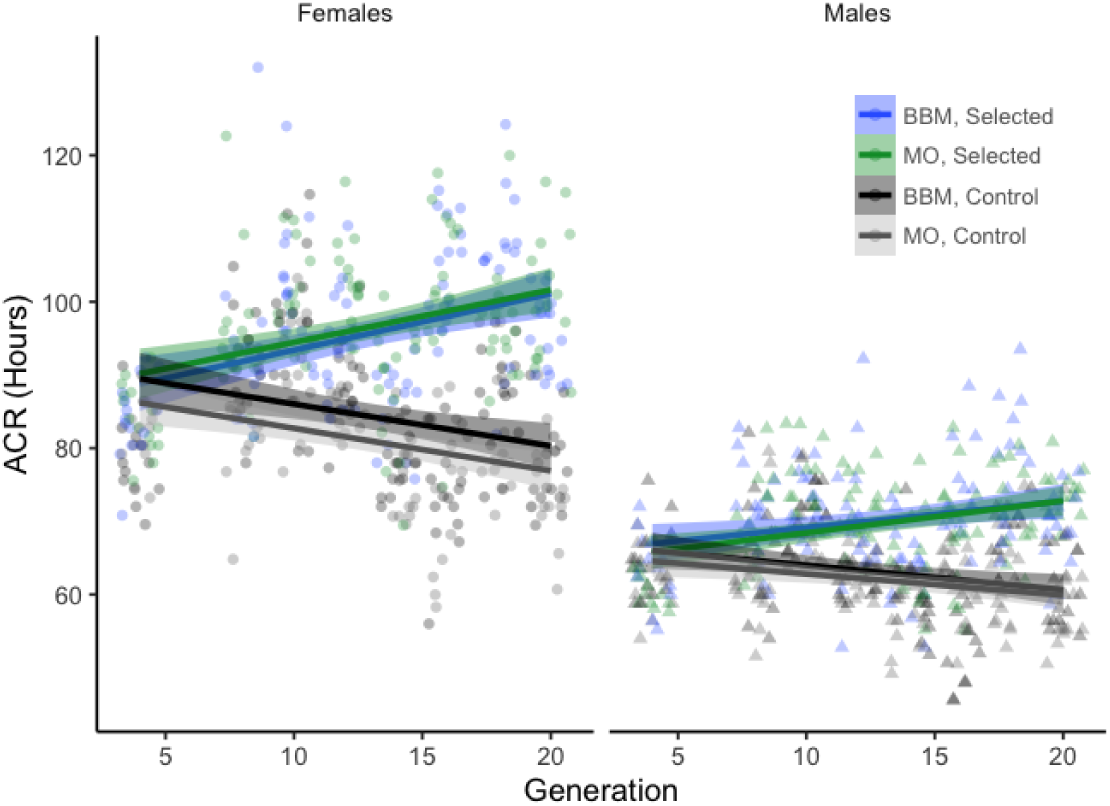
Selection increased adult copper resistance (ACR) in adult flies in both BBM and MO-derived selecte cages (P < 0.00001; Selection x Generation: P < 0.0001). Females responded more strongly to selection compare to males (P < 0.004), and Control cages generally retained baseline levels of ACR, despite a non-significant trend suggesting loss of ACR. Shading describes the 95% CI of the regression. Points indicate mean ACR per vial of 20 flies and symbol shape distinguishes female and male flies. Points are jittered within each generation. ANCOVA results are presented in Table S3.

Both males and females responded to copper selection, though to different degrees (Sex x Selection: F_(1,928)_ =31.81, P < 0.00001). Overall, males had much lower resistance to copper stress (F_(1,928)_ = 1992.26, P < 0.00001; Figure 3), and males displayed a weaker response to copper selection compared to females descended from either collection site (Selection x Sex x Generation: F_(1,928)_ = 8.52, P < 0.004; Figure 3). Sex-specific differences in the response of ACR to selection are likely influenced by the artificial selection process: adults were exposed until approximately 50% of had died, but we expect that surviving females had previously mated to potentially less copper-resistant males. Therefore, it is likely that our selection pressure was more intense for females compared to males. It is also possible that, given males have lower stress resistance, they may also have more limited capacity to improve ACR. Post hoc comparisons of the first and last generations of selection showed that ACR remained essentially constant in the control cages in both sexes (BBM Females G4 vs G20: Bonferroni adj P = 1.0, MO Females G4 vs. G20: adj P = 0.07; BBM Males G4 vs. G20: adj P = 1.0; MO Males G4 vs. G20: adj P = 1.0; Figure 3).

### Adult copper selection maintained resistance to lead and cadmium

Copper stress has been previously shown to influence the expression of genes that also respond to lead and cadmium (38), and we previously identified SNPs near broad categories of genes that respond to oxidative stress, chemical stressors, and mitochondrial function that were associated with copper resistance in wild-derived flies (35). Therefore, we were interested in whether artificial selection for copper resistance would result in a change in resistance to heavy metals that were not the target of selection (lead and cadmium).

Overall, the effect of selection on ADR (cadmium resistance) was significant (Selection: F_(1,616)_ = 70.39, P < 0.00001; Selection x Generation: F_(1,616)_ = 30.12, P < 0.00001; Figure 4A; Table S4). Interestingly, the effect of selection was driven by a loss of ADR in control cages in contrast to increased resistance in selection cages. Female ADR was significantly higher compared to males (Sex: F_(1,616)_ = 1010.28, P < 0.00001), and females from control cages lost ADR over the duration of the selection experiment (BBM Female G4 vs. G20: adj P < 0.007, MO Female G4 vs. G20: adj P < 0.00001). A similar non-significant trend was observed in males from both populations.

**Figure 4.**
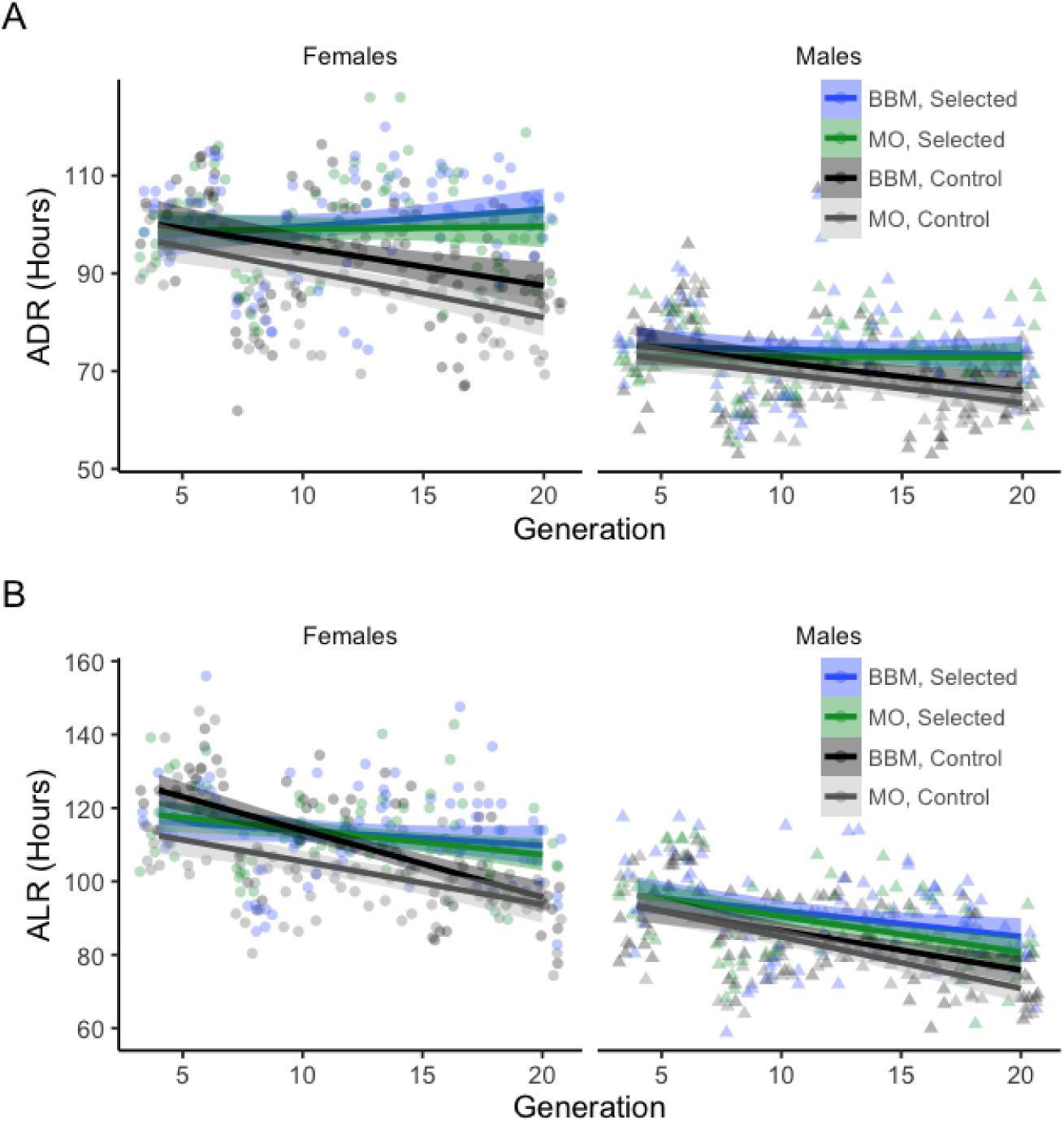
Selection for copper resistance influenced cadmium and lead resistance in adult flies. A. ADR remained constant in response to selection and declined in control cages in females (Selection x Generation: P < 0.00001). B. ALR also did not increase with copper selection but rather remained constant in selected cages (G4 vs. G16 in Selected Cages: Bonferroni adj P = 1.0 for all comparisons). ALR decreased in BBM-derived control cages (G4 vs. G16 in Selected Cages: Bonferroni adj P < 0.001 for males and females), suggesting that copper selection has th potential to maintain ALR. Points indicate mean resistance per vial of 20 flies and symbol shape distinguishes female and male flies. Points are jittered within each generation, and shading indicates the 95% CI of the regression line. ANCOVA results are presented in Table S4.

ALR (adult lead resistance) was also influenced by copper selection, with selection and control cages becoming more distinct over time (Selection: F_(1,616)_ = 52.71, P < 0.00001; Selection x Generation: F_(1,616)_ = 16.87, P < 0.00001; Figure 4B). Unlike ADR, flies from selected cages were more likely to have a slight loss of ALR (BBM Female G4 vs. G20: adj P < 0.005, MO Female G4 vs. G20: adj P = 0.57; BBM Male G4 vs. G20: adj P < 0.0007, MO Male G4 vs. G20: adj P < 0.03). Similar to ADR, females from control cages derived from BBM lost resistance to lead most drastically over the course of the experiment, although the decline was also significant in MO-derived control females (BBM Female G4 vs. G20: adj P < 0.00001, MO Female G4 vs. G20: adj P < 0.00001). Males followed a similar trend with loss of ALR in control cages that reached significance for MO-derived control males (BBM Male G4 vs. G20: adj P = 0.05, MO Male G4 vs. G20: adj P < 0.00001).

Consistent with the lack of increased ADR and ALR in selected cages in response to copper selection, we found that variation in ACR did not explain variation in either trait after accounting for generation and cage (ADR: F_(1,83)_ = 0.24, P = 0.62; ALR: F_(1,77)_ = 0.02, P = 0.88; Pearson Correlation Coefficients < 0.17; Figure 5; Figure S2; Table S5). Taken together, our findings suggest that selection for copper resistance does not result in a correlated increase in lead or cadmium resistance at a phenotypic level that can be detected in our study (Figure S2). However, it remains possible that copper selection could maintain resistance to the non-target heavy metals, as ADR and ALR tended to decrease in cages not subjected to selection (Figure 4B and D). Costs associated with maintaining resistance to chemical stressors warrants further investigation in future studies.

**Figure 5.**
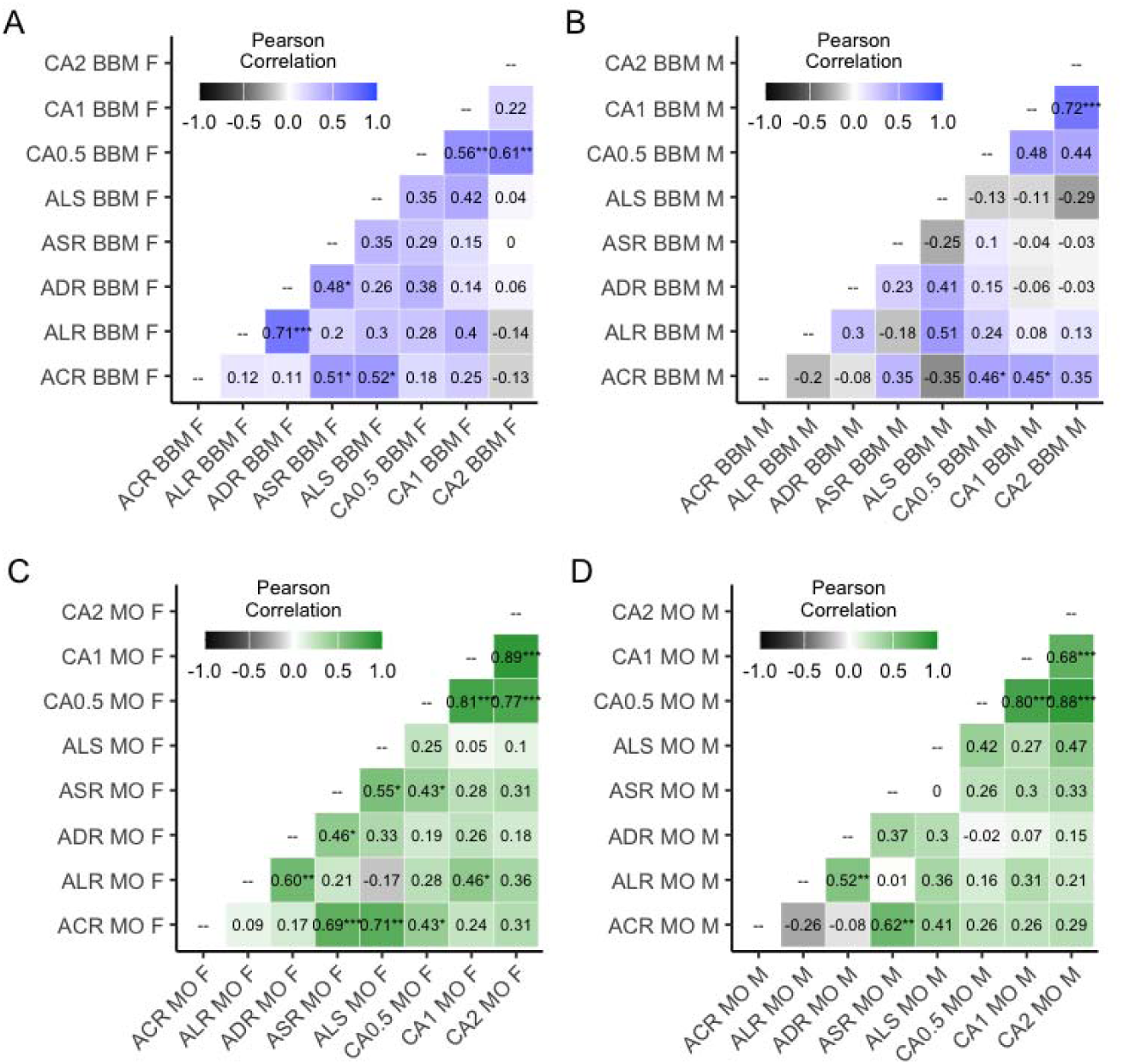
Pearson correlations between each trait. A. and B. present correlation coefficients (r) for BBM female and male estimates. C. and D. present MO female and male estimates. ACR = Adult Copper resistance, ALR = Adult Lead Resistance, ADR = Adult Cadmium Resistance, ASR = Adult Starvation Resistance, ALS = Average Lifespan, CA0.5 = Copper Aversion of 0.5mM CuSO_4_, CA1 = Copper Aversion of 1mM CuSO_4_, CA2 = Copper Aversion of 2mM CuSO_4_. Color scheme differentiates Collection Site (blue = BBM, green = MO). Each axis category indicates Trait, Collection Site (BBM or MO), and Sex (F or M).

### Starvation resistance increased in response to copper selection

Because starvation and copper were initially correlated in each of the collection sites sampled (Figure 1) and because of the potential for off-target selection for starvation resistance through our food-based protocol, we assessed ASR throughout the eight generations of selection in our study. Selection for adult copper resistance had a strong positive effect on starvation resistance (ASR) (Selection: F_(1, 1047)_ = 278.23, P < 0.00001; Selection x Generation: F_(1,1047)_ = 206.41, P < 0.00001; Figure 6; Table S6). The response to selection was slightly stronger in BBM-derived cages compared to MO-derived cages (Collection Site x Generation: F_(1,1047)_ = 26.02, P < 0.00001). ASR was significantly higher in females regardless of collection site (Sex: F_(1,1047)_ = 3921.59, P < 0.00001; Sex x Population: F_(1,1047)_ = 0.03, P = 0.87), and the response to selection was also greater in females compared to males (Sex x Selection: F_(1,1047)_ = 22.66, P < 0.00001). ASR in control cages remained largely constant in both sexes with no significant decline (Figure 6).

**Figure 6.**
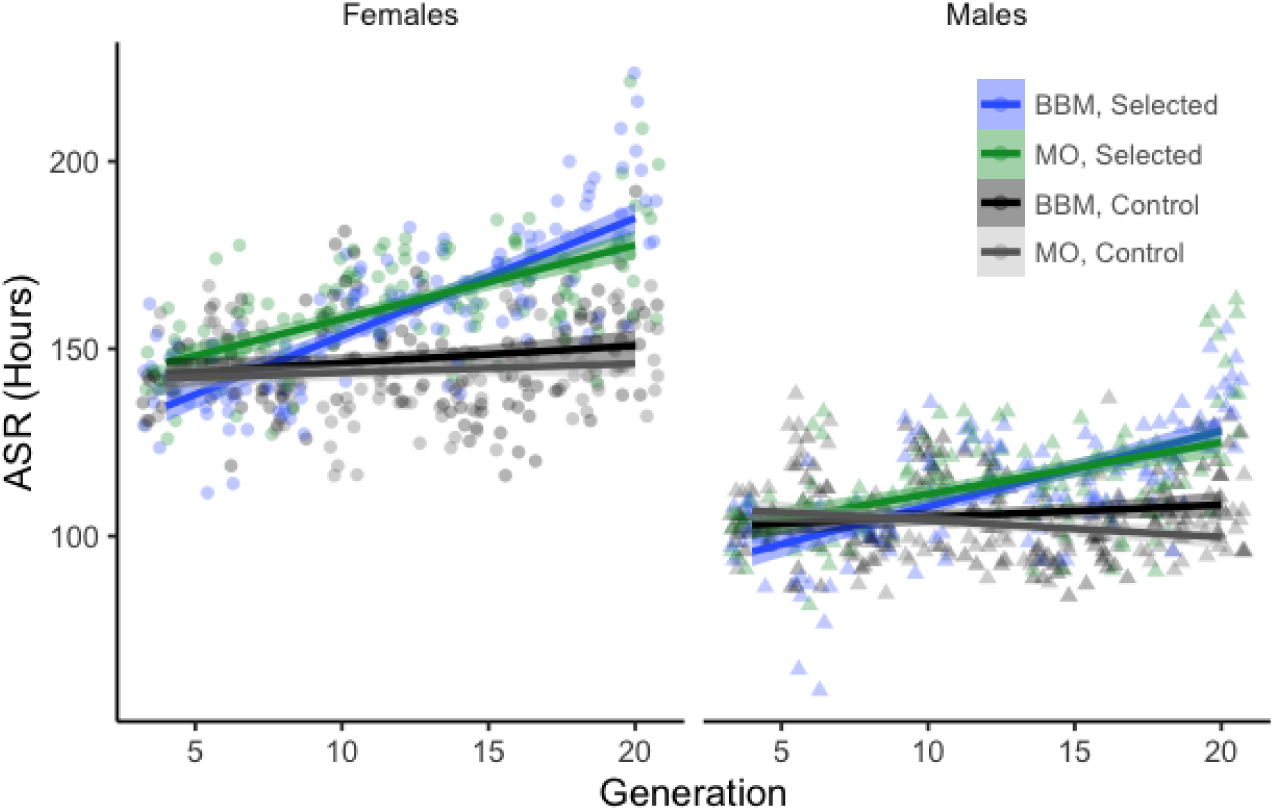
Adult starvation resistance (ASR) increased in response to copper selection in males and females derive from MO and BBM (P < 0.00001). The increase in ASR was slightly greater in flies derived from BBM compared t MO (P < 0.00001). Shading describes the 95% CI of the regression. Control populations retained starvation resistance in both sexes with no statistical support for loss of ASR. Points indicate mean ACR per vial of 20 flies and symbol shape distinguishes female and male flies. Points are jittered within each generation. ANCOVA results ar presented in Table S6.

After accounting for cage and generation, we found that ASR was significantly positively correlated with ACR in the cages that had undergone selection (ACR: F_(1,83)_ = 9.92, P < 0.003; Sex x ACR: F_(1,83)_ = 0.022, P = 0.88; Figure S3; Table S7). The correlation was significant for BBM females and MO males and females (Figure 5). While the correlations are strong and positive, the increase in ACR is not entirely due to an increase in ASR. At most (MO female ACR vs. ASR r = 0.69), variation in ASR accounted for 47.61% of variation in ACR and at least (BBM male ACR vs. ASR r = 0.35) ASR accounted for 12.25% of variance in ACR (Figure 5).

### Behavioral response to copper selection

In addition to altering physiological resistance to a selection pressure, exposure of insects to chemical stressors also has the potential to influence behavior through increased aversion to a toxic chemical or ability to modulate consumption to a level that is sublethal (52,53). We assessed aversion to a low doses of copper (0.5, 1, and 2mM CuSO_4_) in our control and selection cages flies using the Microplate Feeding Assay (58).

In G4, all flies displayed strong aversion to even the lowest concentration of copper tested (0.5mM) (Figure 7; Table S8), and the level of aversion was dose-dependent (Treatment: F_(2,18536)_ = 1599.98, P < 0.00001). Aversion to different concentrations of copper was positively correlated in MO-derived flies and less so in BBM-derived flies from Selection cages (Figure 5). Over time, flies lost the strong aversion to copper to a similar degree across all copper concentrations tested (Treatment x Generation: F_(1,18536)_ = 2.74, P = 0.06). The rate of copper aversion loss was most pronounced in control cages (Selection x Generation: F_(1,18536)_ = 7.20, P < 0.008); however, this difference is not likely biologically relevant given that copper aversion was not different among G20 flies. By G20, flies from all cages were equally likely to consume control food or food with 0.5 mM CuSO_4_. Aversion to copper at any concentration was not correlated with ACR (0.5mM: F_(1,83)_ = 2.60, P = 0.11; 1mM: F_(1,83)_ = 2.32, P = 0.13; 2mM: F_(1,83)_ = 0.34, P = 0.56; Table S9). Thus, we conclude that the loss of copper aversion over generations is not a correlated response to increased ACR. Our data do not provide any evidence that selecting for increased adult copper resistance influences the capacity of flies to detect copper in food or that it increases aversion to copper contaminated food. Flies may lose aversion to copper in food through other mechanisms, such as drift at genes that are not the target of selection for increased physiological copper resistance. Further, one might hypothesize that a correlated response in ASR to copper selection may be driven by increased avoidance to food containing copper in selected cages. Our data do not support this hypothesis and suggest at least partial genetic independence of behavioral and physiological responses to copper stress. This pattern warrants future investigation.

**Figure 7.**
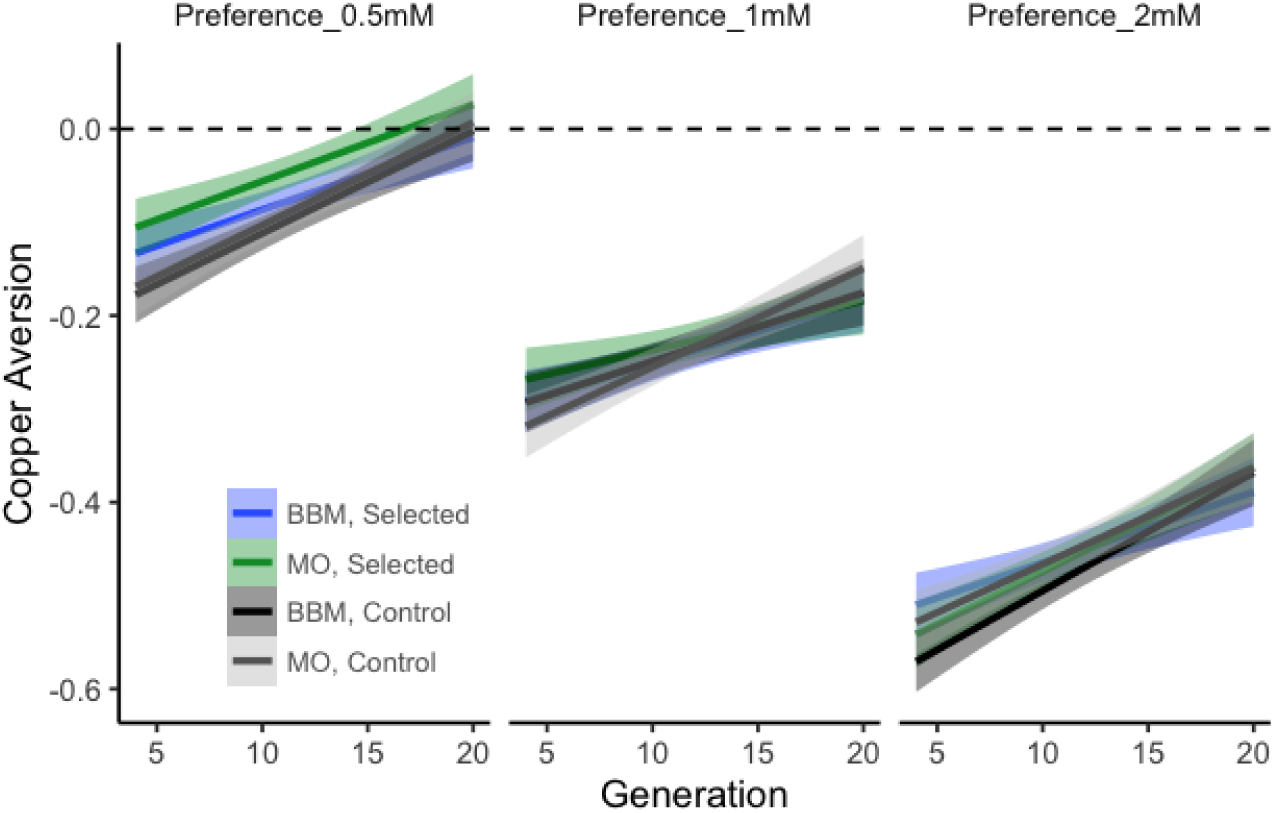
Flies tended to lose aversion to copper regardless of collection site, sex, and selection for adult copper resistance. Shading represents the 95% CI of the regression line shown for visualization and the dashed line highlights the y intercept at 0 and no difference in choice between copper and control food. Negative values indicate preference for control food and aversion to copper, and concentrations tested are indicated in each panel title (e.g. Preference_0.5mM is preference for 0.5mM CuSO_4_). Statistics are presented in Table S8.

### Selection for copper resistance increases lifespan

In addition to stress-response traits, we also measured average adult lifespan under control conditions periodically (G4, G8, G12, G16, G20) during the selection experiment to investigate whether selecting for a chemical stress response trait may influence aspects of life history. We found that selection for copper resistance influenced average lifespan in a collection site-dependent manner (Selection: F_(1,592)_ = 105.29, P < 0.00001; Selection x Generation: F_(1,592)_ = 22.25, P < 0.00001; Selection x Collection Site x Generation: F_(1,592)_ = 13.93, P < 0.0003; Figure 8; Table S10). BBM-derived cages either maintained (in females) or tended to lose longevity (males) in both selected and control cages (Selected cages G4 vs G20: Bonferroni adj P = 0.76; Control cages G4 vs G20: adj P = 1.0), but MO-derived selection cages gained on average 13.07 days of lifespan compared to control cages (Figure 8).

**Figure 8.**
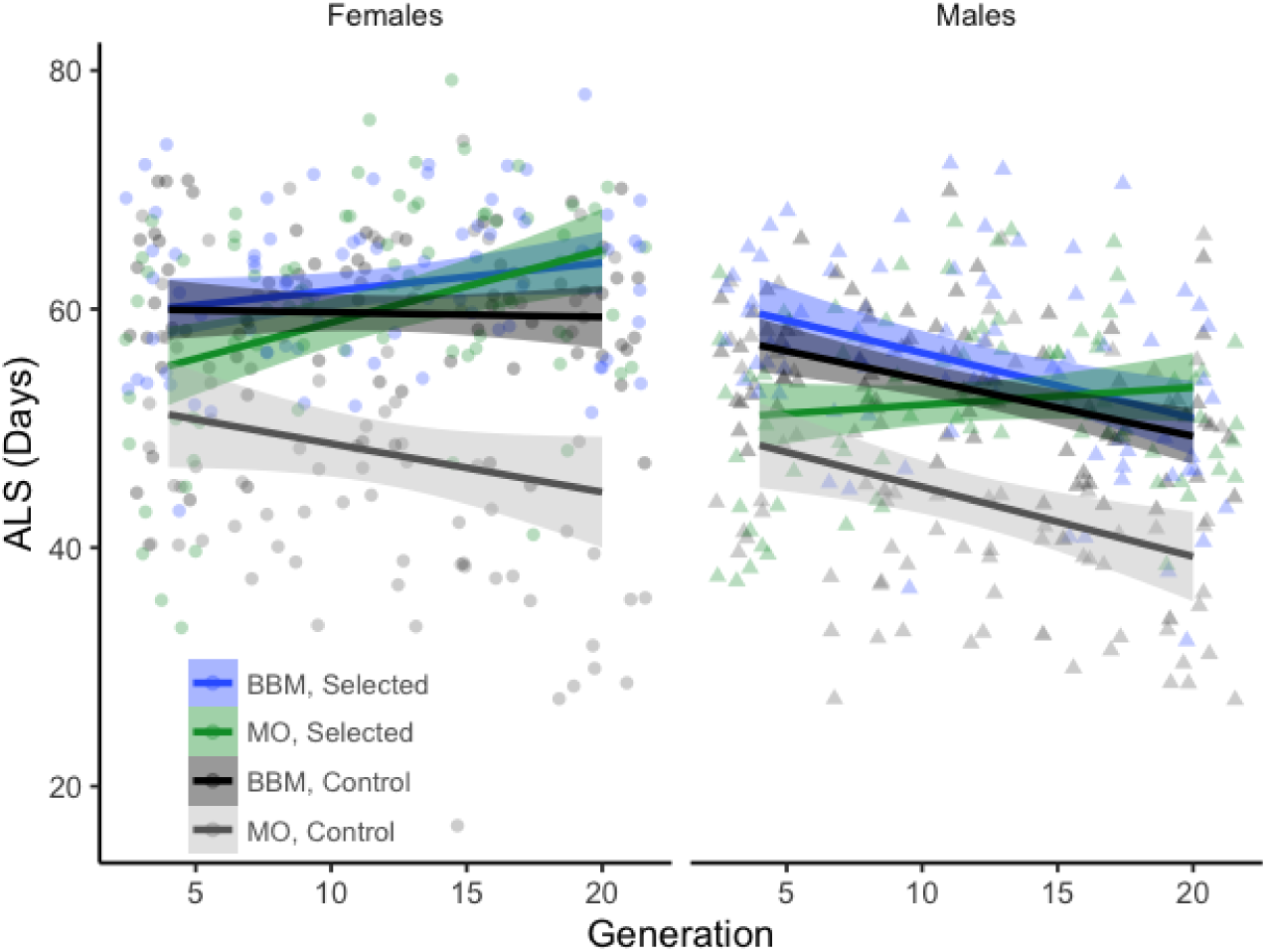
Adult copper selection had population- and sex-specific effects on longevity (ALS). MO-derived selected cages gained longevity overall in both males and females and males from MO control cages lost longevity by G20. BBM-derived cages tended to maintain lifespan throughout selection except for males from selected cages, which lost 10.30 days of average lifespan by G20. Points represent the vial-level response of 20 flies and are jittered. Females and males are distinguished by symbol, and shading represents the 95% CI of the regression line shown for visualization. Statistics are presented in Table S10.

Sex contributed to differences in average lifespan, with females living on average 6.12 days longer than males (Sex: F_(1,592)_ = 106.32, P < 0.00001; Figure 8). We detected a significant interaction between sex and generation (F_(1,592)_ = 20.30, P < 0.00001) that was in part driven by a stronger response to selection in MO-derived females compared to MO-derived males (Selected MO Females G4 vs G20: 10.6 day increase, Bonferroni adj P < 0.002; Selected MO Males G4 vs G20: 0.70 day increase, adj P < 0.003). MO-derived males from control cages lost longevity (Control MO Males G4 vs G20: adj P < 0.0007). MO-derived control females followed the same non-significant trend (Control MO Females G4 vs G20: adj P = 0.06). In BBM-derived cages, females did not appear to respond to selection and maintained longevity in control cages (all G4 vs G20 adj P = 1.0). However, BBM-derived males tended to lose longevity in both selected and control cages, although this trend only reached significance in males from selected cages (Control G4 vs G20: adj P = 0.26; Selected G4 vs G20: 10.30-day decrease, adj P < 0.003).

Consistent with patterns observed in Figure 8, we found that ACR and ALS were correlated and that the correlation depended upon collection site in selected cages after accounting for cage and generation (ACR: F_(1,47)_ = 12.01, P < 0.002; ACR x Collection Site: F_(1,47)_ = 5.60, P < 0.03; Figure S5; Table S11). The correlation between ACR and ALS was strongest in MO females with variation in ACR explaining 50.4% of variation in ALS (Figure 5) followed by BBM females (27.04% variation in ALS explained; Figure 5). The correlations between ACR and ALS were not significant in male flies from selected cages, although they followed a trend similar to that demonstrated in Figure 8.

## Discussion

### Copper resistance increased in flies from two ecologically distinct collection sites

We collected flies from two sites chosen for their proximity to the Copper Basin, USA to explore the effects of artificial selection for copper resistance on target and off-target stress resistance, fitness, and behavioral traits. We previously demonstrated that copper resistance is high in the BBM population compared to other sites outside the Copper Basin despite the copper mine being retired for several decades (35). In contrast, MO, a large (27-acre) orchard, is likely influenced by a combination of historical proximity to copper mines (being 16.84km southeast of BBM) as well as contemporary routine use of organophosphate and Spinosad chemical control of pests (personal communication to ERE from MO owner). As many insecticides including those used at MO result in oxidative stress (15,64), we expected copper resistance to be high in MO as well and were interested in differences in the response to selection between the two collection sites given these differences in anthropogenic impact.

From the start of our study, we found that ACR in flies from MO was comparable if slightly lower than that of high resistance BBM-derived flies (Figure 1). Correlations between resistance to heavy metals and insecticides have been previously reported in several species (reviewed in 65) including mosquitoes (15,66), the beet armyworm (9), and plant hoppers (13). Further, some pesticides contain copper as a means of control (67) and adaptive responses to copper-containing pesticides have been documented in fungi (68,69) and bacteria (70). Consistent with previous reports (7,35), our sampling of flies from the active agricultural site suggests that insecticide use is associated with increased resistance of *D. melanogaster* to copper toxicity as well.

Prior exposure to insecticides may also have influenced the adaptive potential of MO-derived flies versus those collected from BBM. Although the response to selection was statistically indistinguishable between the replicated selection cages (Figure 3; Table S3), MO-derived flies experienced a larger overall change in ACR compared to BBM. Comparison of flies as close to wild-caught as possible (G2) and G20 revealed that the increase in ACR was greater in MO males and females compared to BBM flies (F_(1,739)_ = 7.79, P < 0.006; Figure 9). In addition, our replicate cages demonstrated that the response to selection from both collection sites was largely consistent within sex and collection site except for in females from MO selected cages (Tukey HSD adj P > 0.05 except female MA2 vs MA3: adj P < 0.05). The number of founding females differed between MO (355 females) and BBM (191 females) cages, and increased genetic diversity may have influenced the greater gain in ACR of MO-derived flies.

**Figure 9.**
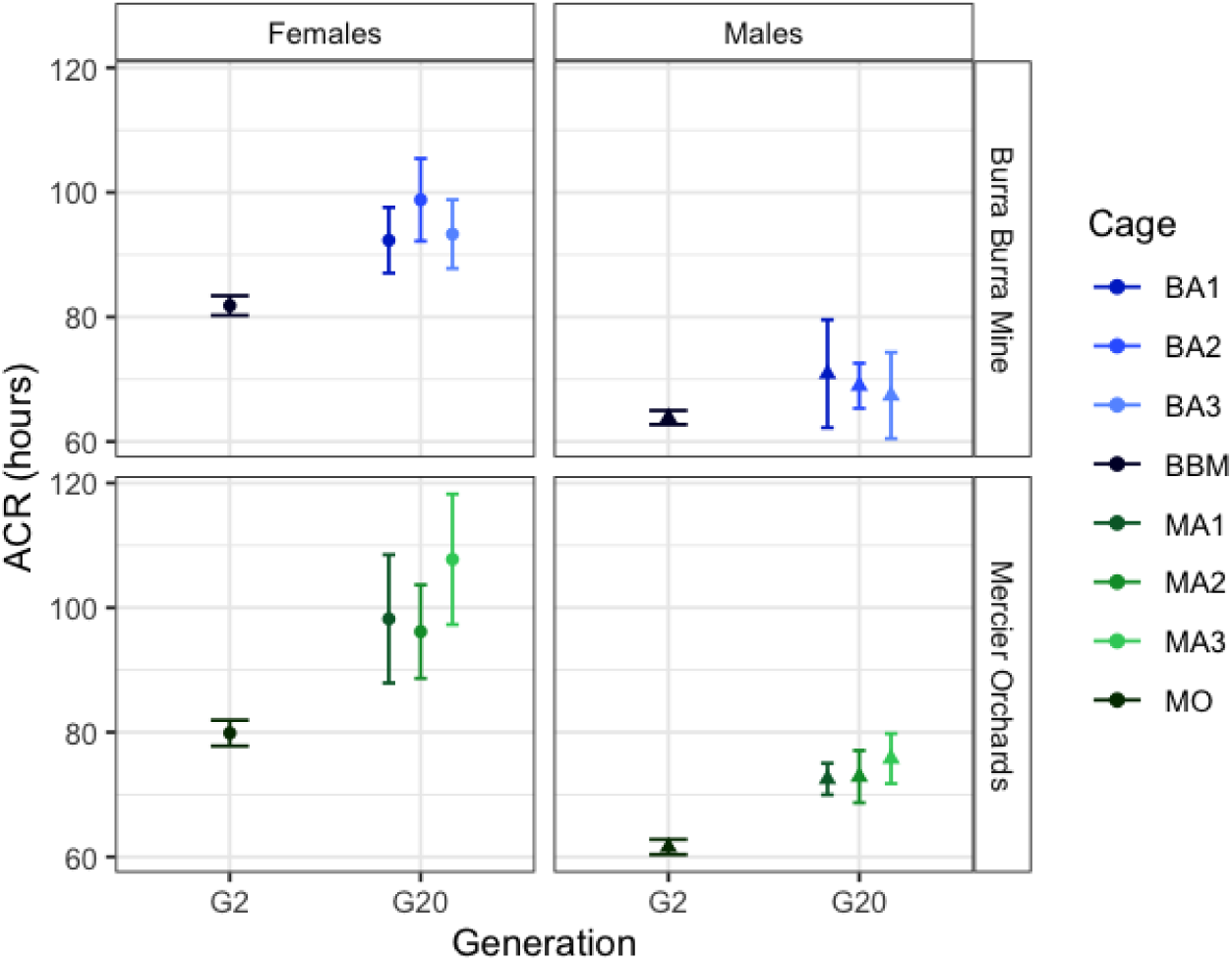
Replicate cages subjected to adult selection for copper resistance responded similarly within sex and collection site. The overall response to copper selection was greater in male and female flies derived from MO. Dat shown are mean +/-95% CI. Shape indicates sex.

Alternatively, repeated exposure of the MO wild population to chemicals used in pest control may have provided greater genetic predisposition or flexibility to respond to the copper stress used in our study. The potential for pre-adaptation following exposure to a variable, stressful environment may increase the adaptive potential of populations to future novel stressors (10), especially if the genes involved in resistance to current stressors have pleiotropic effects. The two pesticides routinely used at MO fit this description. Spinosad is derived from the bacterium *Saccharophlyspora spinosa* and targets nicotinic acetylcholine receptors in neurological tissue, specifically leading to lysosome dysfunction and accumulation of reactive oxygen species (ROS) that ultimately cause oxidative stress in *D. melanogaster* (71). Organophosphates similarly disrupt neurological function through interaction with acetylcholinesterase, also ultimately leading to increased oxidative stress (72). The oxidative stress response itself is quite broad, including heat shock proteins (HSPs), cytochrome p450s (cyp450s), and glutathione-s-transferases (GSTs) (73–77), all of which have also been demonstrated to respond to oxidative stress induced by heavy metal exposure (50,78–80). The highly polygenic nature of the oxidative stress response and overlap of both mechanism and genes with heavy metal toxicity response may, in part, account for the increase in resistance of MO-derived selection cages. Future examination of allelic variants underlying the shift in ACR between and among the MO and BBM cages will assist with determining whether these broad oxidative stress response gene families are contributing to copper adaptation and whether the relatively consistent response to selection across different cage replicate is also repeatable at the genetic level.

### Selection for copper resistance has the potential to influence resistance to other stressors

Selection for increased resistance to copper has the potential to result in correlated responses to other metals. This potential stems from pleiotropic effects of genes involved in metal metabolism, such as metal scavenger proteins (metallothioneins) and metal-dependent or oxidative stress-induced transcription factors (e.g. MTF-1, CncC/Nrf2, AP-1), which have been linked to stress induced by copper, cadmium, and lead (31,37,38,41,81).

A recent study in *Saccharomyces cerevisiae* examined the evolution of cross-tolerance to several heavy metals including copper and cadmium (12). They found that cross-tolerance can evolve, but that it is difficult to predict and dependent upon the nature of the genetic changes that have led to increased resistance. In particular, while all strains artificially selected for increased copper resistance also had elevated resistance to manganese, only a subset had increased cadmium resistance despite similarities in metal properties and a large overlap in genes responsive to cadmium and copper (12). Our examination of cross-tolerance between copper, cadmium, and lead revealed a similarly complex relationship. Copper selection did not result in a correlated increase in resistance to either non-essential metal (Figure 4; Figure S2); however, cages subjected to copper selection retained resistance to both cadmium and lead (Figure 4). Cages maintained under control conditions significantly declined in cadmium and lead resistance over time, suggesting that there may be energetic costs associated with maintaining resistance to non-essential heavy metals.

In contrast to the lack of increased off-target metal resistance, we observed a collection site-specific, strong correlated response to starvation in selected cages (Figure 6). This association between copper resistance and starvation resistance has been previously reported in *D. melanogaster* (35), and starvation resistance has also been previously linked to increased paraquat resistance (82). The correlation between starvation and copper resistance may be related to reductions in metabolic rate and activity levels as a result of selection (57,83,84). Response to metal toxicity is energetically taxing, and disruption to copper homeostasis can negatively impact the typical role of copper ions in mitochondrial function and metabolism (30). For example, in the ragworm (*Nereis diversicolor*), resistance to copper was linked to reduced growth and reproduction as well as lipid and carbohydrate reserves (85). Similar energetic costs were observed in a freshwater snail (*Planorbella pilsbyri*) (86). The increase in starvation resistance observed in both MO and BBM-derived selected flies may be due adaptive responses to energetic stress.

Behavioral resistance my also contribute to the mechanism underlying the link between copper and starvation resistance. In addition to altering physiological resistance to a selection pressure, exposure of insects to chemical stressors can increase aversion to a toxic chemical or the ability to modulate consumption to a level that is sublethal (52,53). We previously found that *D. melanogaster* avoid copper-contaminated food and that more copper sensitive strains may consume more copper within a 24-hour period (33). If selection for copper resistance results in increased aversion to copper, it is conceivable that both copper resistance and starvation resistance may increase because flies avoid eating copper-contaminated food for as long as possible. However, our study did not provide any evidence that selecting for increased adult copper resistance influences the capacity of flies to detect copper in food or that it increases aversion of copper contaminated food (Figure 5; Figure 7). In fact, flies from all cages (Control and Selection) gradually lost aversion to copper over time, suggesting genetic independence of behavioral and physiological responses to copper selection in the populations examined in our study.

### Fitness effects of copper selection are population specific

Negative consequences of metal exposure on survival and health have been extensively demonstrated in fish, arthropods, plants, and humans (46,85,e.g. 87,88). More limited attention has been given to correlated shifts in fitness-relevant traits as a result of adaptation to metal stress. Previous work in *D. melanogaster* has found evidence of a trade-off between metal resistance and longevity (51), although others have found no relationship between starvation resistance and lifespan (82). Our study offers a novel perspective on the consequences of adaptation to copper stress by tracking longevity throughout our selection experiment. We found that the effect of copper selection was collection site-specific: BBM flies either maintained or tended to lose longevity independent of selection, while MO-derived flies gained longevity in tandem with copper resistance (Figure 8). The mechanism underlying this collection site-specific response requires further investigation, but may be related to shifts in allele frequencies for genes that can influence both traits or (or in addition to) ecological differences between the BBM and MO sites. For example, in *Caenorhabditis elegans*, four genes with pleiotropic effects on metal resistance and longevity have been characterized (89,90). Future analysis of the genetic basis of copper adaptation in the MO and BBM populations will allow us to investigate the potential for allele frequency shift at genes that influence copper resistance and lifespan.

In addition to possible genetic overlap, resource availability differs drastically between the BBM and MO Collection Sites. Large fruit crops are consistently available as a food source throughout the growing season at MO, whereas BBM lacks cultivated crops with the exception of a single apple tree. Natural selection for higher baseline lifespan in the BBM population may have resulted from more limited or more inconsistent opportunities for oviposition at this site. MO flies that are routinely exposed to insecticides may not be able to survive beyond early adulthood, effectively selecting against long life and favoring flies that reproduce early (91,92). Standing genetic variation in the MO population that overlaps with or is in linkage disequilibrium with variants associated with copper resistance may have increased in frequency during artificial selection, leading to increased lifespan.

## Conclusions

Understanding the consequences of adaptive responses from the perspective of cross-tolerance with other forms of stress can support a broader understanding for how genetically complex traits shift over time. We also gain a deeper understanding of the evolutionary potential of organisms to respond to ever-changing environments. Future genomic exploration and characterization of the traits we characterized in our study will additionally clarify the genetic response to selection. Together, our results underscore the complexity of adaptive shift in polygenic traits and provide a basis for further exploration of costs and correlative change following copper selection.

## List of abbreviations

BBM: Burra Burra Mine
MO: Mercier Orchards
ACR: Adult Copper Resistance
ADR: Adult Cadmium Resistance
ALR: Adult Lead Resistance
ASR: Adult Starvation Resistance
ALS: Average Lifespan
CA(0.5,1,2): Copper Aversion at 0.5, 1, 2mM CuSO_4_

## Competing interests

The authors declare that they have no competing interests.

## Funding

This work was supported by an R00 (NIEHS R00ES033257) awarded to ERE and by the University of Oklahoma.

## Authors’ contributions

ERE conceived and designed the study, carried out artificial selection, and phenotyped ACR, ADR, ALR, ASR, and CA. KA prepared media for all experiments and assisted with data collection. KA also carried out the ALS phenotyping and data preparation. ERE analyzed the data and wrote the first draft of the manuscript. KA and ERE edited the manuscript. All authors read and approved the final manuscript.

## Acknowledgments

The Burra Burra Mine and surrounding area was originally part of the Cherokee Nation territory before their forced removal by the United States in 1836. We acknowledge, honor, and respect the diverse indigenous peoples connected to this land. We thank The Ducktown Basin Museum and Mercier Orchards for allowing us access to sites to collect flies.

---

**Figure S1.**
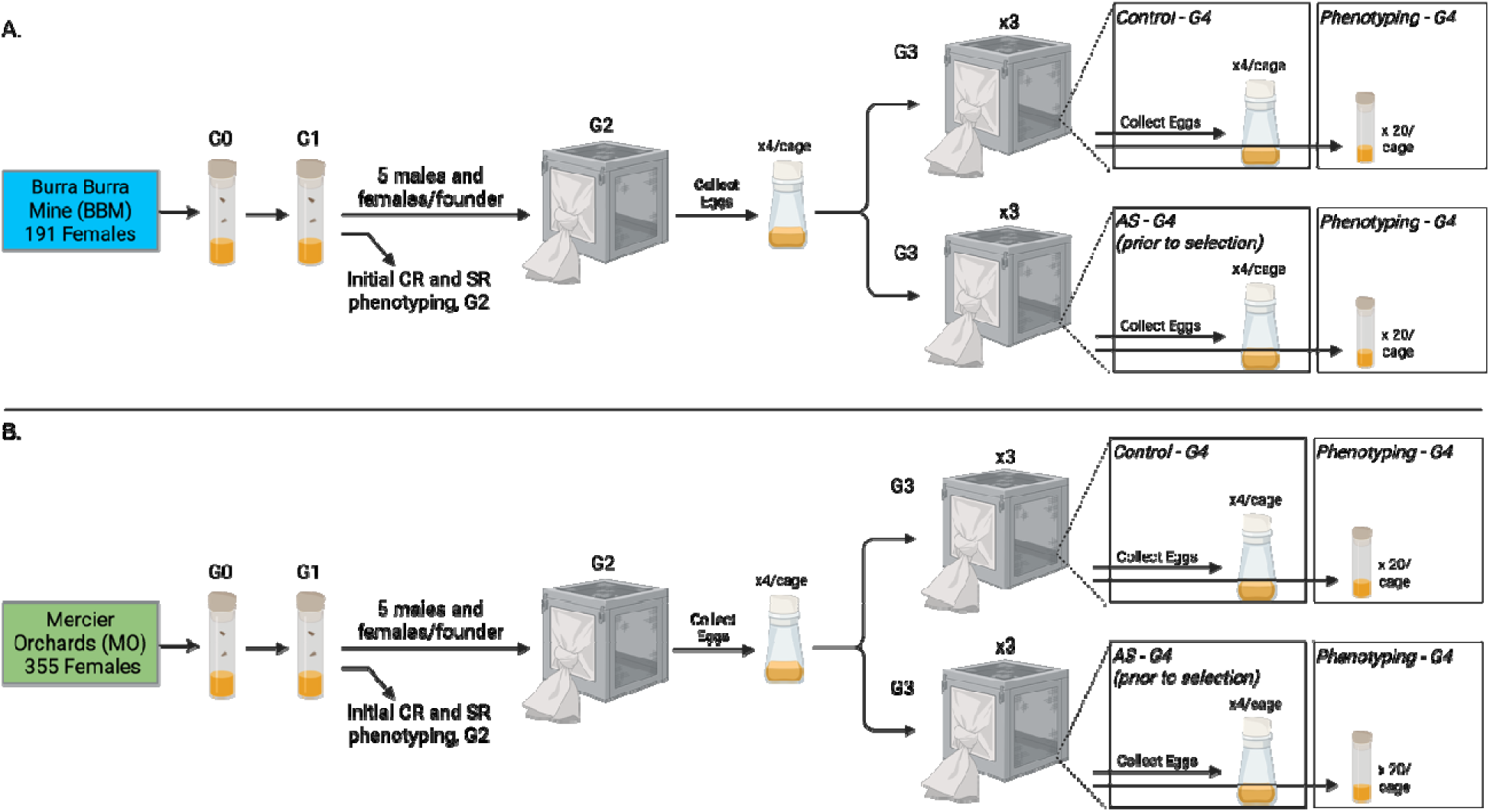
Replicated cages were established from two wild populations of D. melanogaster. A. We verified 191 gravid BBM females were D. melanogaster and used their offspring to establish one Generation 2 (G2) population cage and to measure baseline levels of copper resistance (CR) and starvation resistance (SR). After one generation of interbreeding, eggs were collected from G2 cages to establish three control and three adult selection (AS) G3 cages. Following a second generation of interbreeding, eggs were collected to establish G4 cages. We additionally collected eggs in vials to measure the following traits in G4 individuals: Adult Copper Resistance (ACR), Adult Lead Resistance (ALR), Adult Cadmium Resistance (ADR), Adult Starvation Resistance (ASR), Copper Aversion (CA), and Adult Lifespan (ALS). B. The same steps were followed to establish MO cages from the 355 species-verified founder females. This figure was created with BioRender.com.

**Figure S2.**
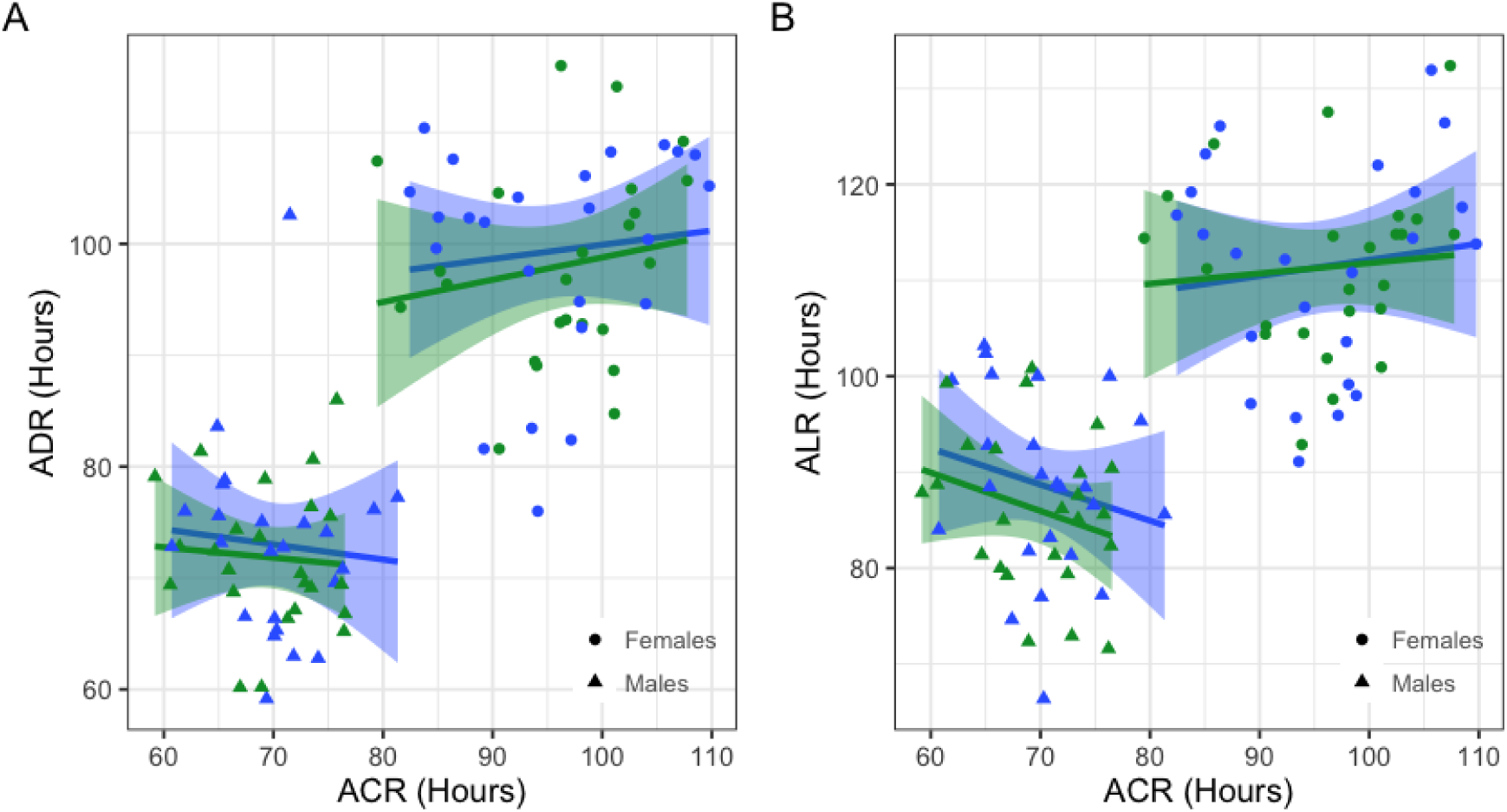
Copper resistance (ACR) is not correlated with resistance to non-target heavy metals. A. Variation in ADR (cadmium) is not explained by variation in ACR (P = 0.62), nor was variation in ALR (lead) (B. P = 0.88). In both plots, points indicate sex-specific averages for each of the Adult Selection cages. All generations are plotted together. Blu represents BBM; green represents MO. Shading indicates the 95% CI of linear regressions used for visualization. Note that y axes are not consistent between A and B. Results from each ANCOVA are presented in Table S5.

**Figure S3.**
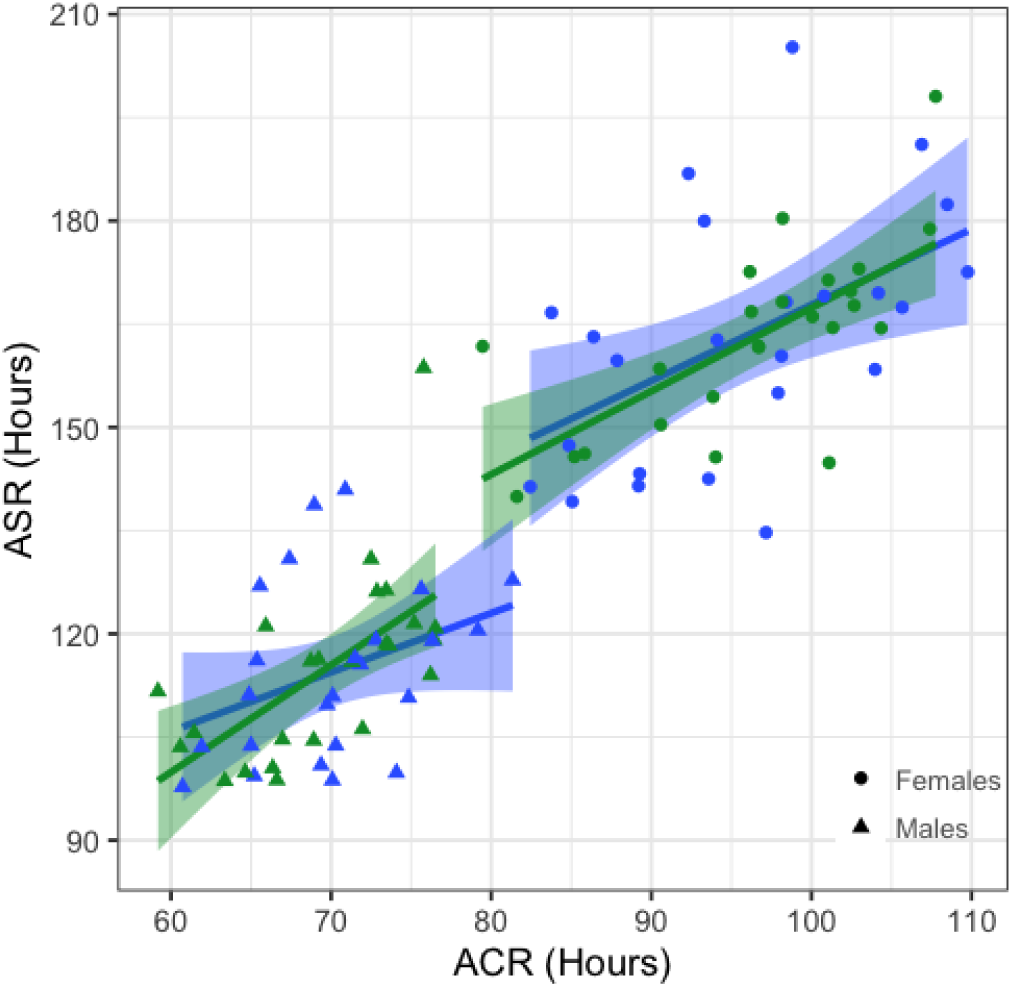
ACR and ASR were positively correlated, accounting for cage and generation (P < 0.003). Points indicate sex-specific averages for each of the Adult Selection cages. Blue represents BBM; green represents MO. All generations are plotted together. Shading indicates the 95% CI of a linear regression used for visualization. Statistic are reported in Table S7.

**Figure S4.**
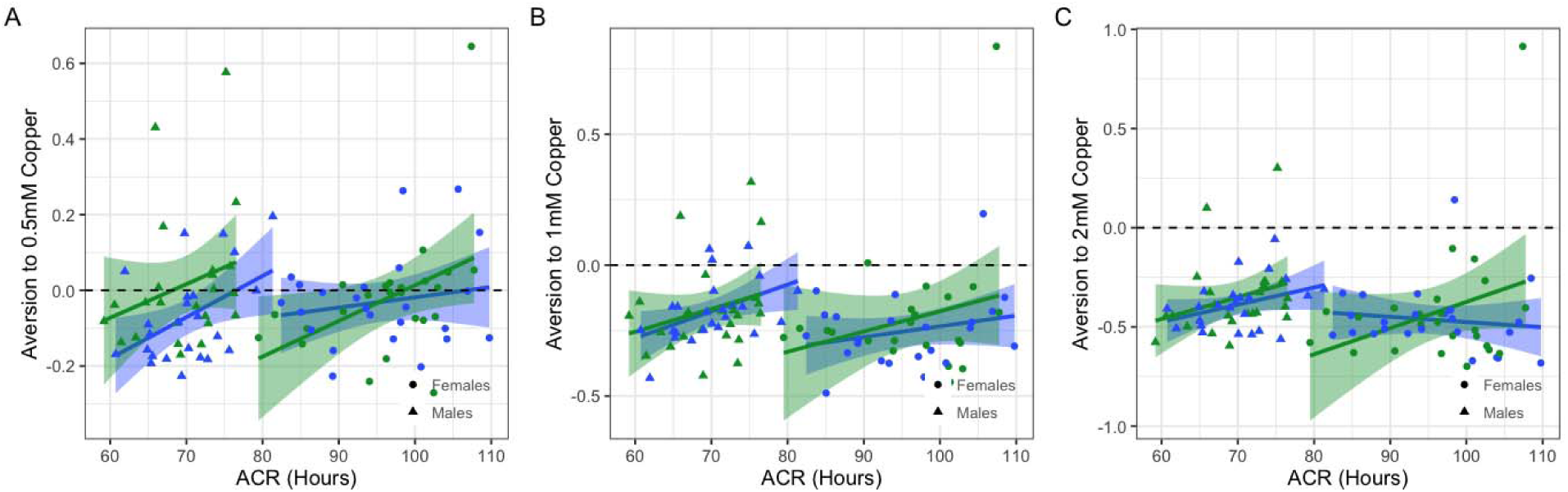
After accounting for Cage and Generation, copper aversion was not correlated with ACR at 0.5, 1, or 2mM CuSO_4_. In each plot, negative values indicate preference for control food and aversion to copper. Blue represents BBM-derived cages; green represents MO-derived cages. Points are average estimates per Cage, and all generations are shown. Symbol indicates sex. Shading represents the 95% CI of the regression shown for visualization Table S9.

**Figure S5.**
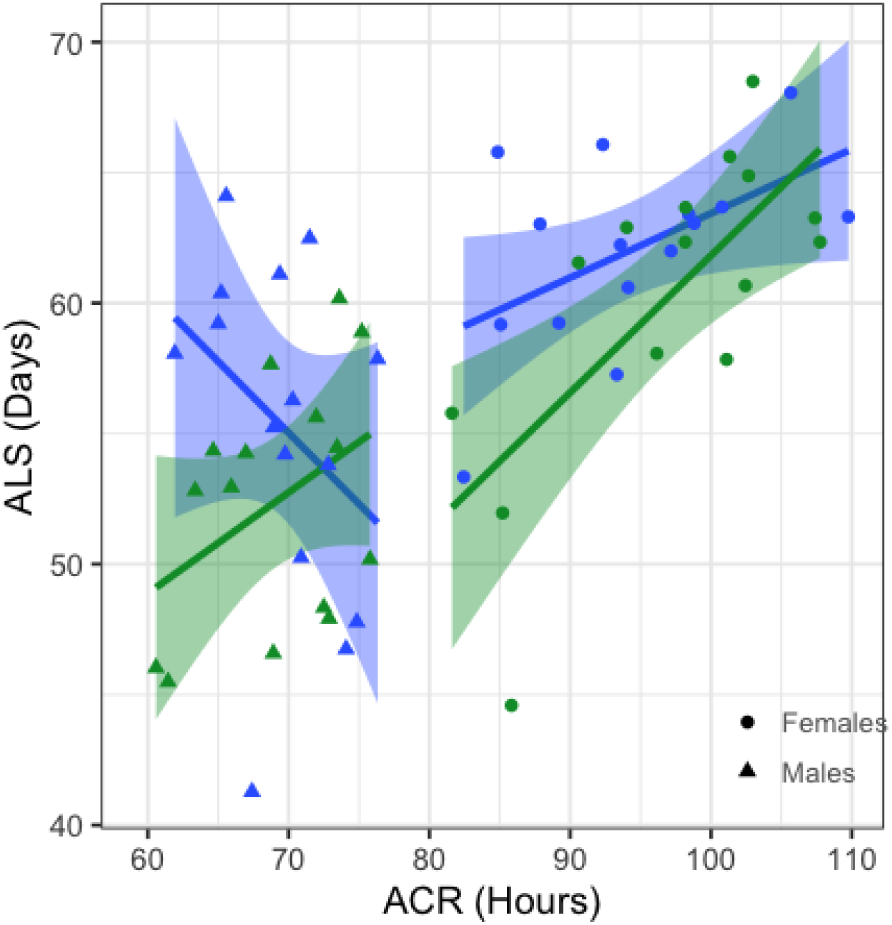
ACR and ALS were positively correlated in females derived from BBM (blue) and MO (green) in Selected cages after accounting for Cage and Generation (P < 0.002). Male ACR was not correlated with ALS. Points indicate average responses for each Selected cage and all generations are presented. Plot symbols distinguish sex. Shading represents the 95% CI of the regression. Statistics are presented in Table S11.

**Table S1.** ANOVAs for ACR and ASR in G2 wild-derived flies.

| Effect | DF | SS | MS | F Value | P Value |
| --- | --- | --- | --- | --- | --- |
| **Adult Copper Resistance (ACR)** |  |  |  |  |  |
| Sex | 1 | 56198 | 56198 | 553.29 | < 0.00001 |
| Collection Site | 1 | 777 | 777 | 7.65 | < 0.006 |
| Sex x Collection Site | 1 | 6 | 6 | 0.06 | 0.81 |
| Residuals | 679 | 68966 | 102 |  |  |
| **Adult Starvation Resistance (ASR)** |  |  |  |  |  |
| Sex | 1 | 208490 | 208490 | 793.61 | < 0.00001 |
| Collection Site | 1 | 1232 | 1232 | 4.69 | < 0.04 |
| Sex x Collection Site | 1 | 271 | 271 | 1.03 | 0.31 |
| Residuals | 591 | 155263 | 263 |  |  |

**Table S2.** ANOVA for ACR and ASR in G2 wild-derived flies versus G4 cages prior to selection.

| Effect | DF | SS | MS | F Value | P Value |
| --- | --- | --- | --- | --- | --- |
| **Adult Copper Resistance (ACR)** |  |  |  |  |  |
| Sex | 1 | 67491.61 | 67491.61 | 738.06 | < 0.00001 |
| Collection Site | 1 | 688.06 | 688.06 | 7.52 | < 0.007 |
| Cage* | 12 | 622.03 | 51.84 | 0.57 | 0.87 |
| Sex x Collection Site | 1 | 28.94 | 28.94 | 0.32 | 0.57 |
| Sex x Cage* | 12 | 403.79 | 33.65 | 0.37 | 0.97 |
| Residuals | 779 | 71235.15 | 91.44 |  |  |
| **Adult Starvation Resistance (ASR)** |  |  |  |  |  |
| Sex | 1 | 256443.93 | 256443.93 | 1099.10 | < 0.00001 |
| Collection Site | 1 | 864.12 | 864.12 | 3.70 | 0.05 |
| Cage* | 12 | 878.53 | 73.21 | 0.31 | 0.99 |
| Sex x Collection Site | 1 | 262.09 | 262.09 | 1.12 | 0.29 |
| Sex x Cage* | 12 | 445.70 | 37.14 | 0.16 | 1.00 |
| Residuals | 691 | 161225.49 | 233.32 |  |  |

\*Cage includes G2 vials and G4 cages

**Table S3.** ANCOVA of the effect of selection on ACR.

| Effect | DF | SS | MS | F Value | P Value |
| --- | --- | --- | --- | --- | --- |
| **Adult Copper Resistance (ACR)** |  |  |  |  |  |
| Sex | 1 | 128807.57 | 128807.57 | 1992.26 | < 0.00001 |
| Selection | 1 | 25302.41 | 25302.41 | 391.35 | < 0.00001 |
| Collection Site | 1 | 237.17 | 237.17 | 3.67 | 0.06 |
| Cage | 9 | 2040.75 | 226.75 | 3.51 | < 0.0003 |
| Generation | 1 | 97.27 | 97.27 | 1.50 | 0.22 |
| Sex x Selection | 1 | 2056.59 | 2056.59 | 31.81 | < 0.00001 |
| Sex x Collection Site | 1 | 10.91 | 10.91 | 0.17 | 0.68 |
| Sex x Cage | 9 | 1043.02 | 115.89 | 1.79 | 0.07 |
| Collection Site x Generation | 1 | 1.11 | 1.11 | 0.02 | 0.90 |
| Selection x Generation | 1 | 6186.88 | 6186.88 | 95.69 | < 0.00001 |
| Sex x Generation | 1 | 9.17 | 9.17 | 0.14 | 0.71 |
| Selection x Collection Site x Generation | 1 | 0.07 | 0.07 | 0.00 | 0.97 |
| Sex x Collection Site x Generation | 1 | 11.92 | 11.92 | 0.18 | 0.67 |
| Sex x Selection x Generation | 1 | 551.15 | 551.15 | 8.52 | < 0.004 |
| Sex x Selection x Collection Site x Generation | 1 | 2.12 | 2.12 | 0.03 | 0.86 |
| Residuals | 928 | 59998.76 | 64.65 |  |  |

**Table S4.** ANCOVAs of the effect of selection on ADR and ALR.

| Effect | DF | SS | MS | F Value | P Value |
| --- | --- | --- | --- | --- | --- |
| **Adult Cadmium Resistance (ADR)** |  |  |  |  |  |
| Sex | 1 | 93047.97 | 93047.97 | 1010.28 | < 0.00001 |
| Selection | 1 | 6483.36 | 6483.36 | 70.39 | < 0.00001 |
| Collection Site | 1 | 964.50 | 964.50 | 10.47 | < 0.002 |
| Cage | 9 | 737.26 | 81.92 | 0.89 | 0.53 |
| Generation | 1 | 1916.94 | 1916.94 | 20.81 | < 0.00001 |
| Sex x Selection | 1 | 781.43 | 781.43 | 8.48 | < 0.004 |
| Sex x Collection Site | 1 | 68.33 | 68.33 | 0.74 | 0.39 |
| Sex x Cage | 9 | 1121.07 | 124.56 | 1.35 | 0.21 |
| Collection Site x Generation | 1 | 43.76 | 43.76 | 0.48 | 0.49 |
| Selection x Generation | 1 | 2773.80 | 2773.80 | 30.12 | < 0.00001 |
| Sex x Generation | 1 | 0.39 | 0.39 | 0.00 | 0.95 |
| Selection x Collection Site x Generation | 1 | 0.08 | 0.08 | 0.00 | 0.98 |
| Sex x Collection Site x Generation | 1 | 80.56 | 80.56 | 0.87 | 0.35 |
| Sex x Selection x Generation | 1 | 316.34 | 316.34 | 3.43 | 0.06 |
| Sex x Selection x Collection Site x<br>Generation | 1 | 12.43 | 12.43 | 0.13 | 0.71 |
| Residuals | 616 | 56734.16 | 92.10 |  |  |
| **Adult Lead Resistance (ALR)** |  |  |  |  |  |
| Sex | 1 | 88365.55 | 88365.55 | 743.00 | < 0.00001 |
| Selection | 1 | 6268.77 | 6268.77 | 52.71 | < 0.00001 |
| Collection Site | 1 | 1462.00 | 1462.00 | 12.29 | < 0.0005 |
| Cage | 9 | 5734.26 | 637.14 | 5.36 | < 0.00001 |
| Generation | 1 | 18270.97 | 18270.97 | 153.63 | < 0.00001 |
| Sex x Selection | 1 | 0.39 | 0.39 | 0.00 | 0.95 |
| Sex x Collection Site | 1 | 96.93 | 96.93 | 0.82 | 0.37 |
| Sex x Cage | 9 | 981.79 | 109.09 | 0.92 | 0.51 |
| Collection Site x Generation | 1 | 13.28 | 13.28 | 0.11 | 0.74 |
| Selection x Generation | 1 | 2006.68 | 2006.68 | 16.87 | < 0.00005 |
| Sex x Generation | 1 | 1.84 | 1.84 | 0.02 | 0.90 |
| Selection x Collection Site x Generation | 1 | 197.76 | 197.76 | 1.66 | 0.20 |
| Sex x Collection Site x Generation | 1 | 264.39 | 264.39 | 2.22 | 0.14 |
| Sex x Selection x Generation | 1 | 329.99 | 329.99 | 2.77 | 0.10 |
| Sex x Selection x Collection Site x<br>Generation | 1 | 251.03 | 251.03 | 2.11 | 0.15 |
| Residuals | 616 | 73261.54 | 118.93 |  |  |

**Table S5.** ANCOVA testing the correlation between ACR and non-target metals.

| Effect | DF | SS | MS | F Value | P Value |
| --- | --- | --- | --- | --- | --- |
| **ACR vs ADR** |  |  |  |  |  |
| Generation | 1 | 333.63 | 333.63 | 4.51 | < 0.05 |
| Cage | 5 | 122.75 | 24.55 | 0.33 | 0.89 |
| Sex | 1 | 16577.86 | 16577.86 | 224.17 | < 0.00001 |
| ACR | 1 | 18.30 | 18.30 | 0.25 | 0.62 |
| Sex x ACR | 1 | 74.03 | 74.03 | 1.00 | 0.32 |
| Sex x Collection Site | 1 | 0.00 | 0.00 | 0.00 | 1.00 |
| ACR x Collection Site | 1 | 1.40 | 1.40 | 0.02 | 0.89 |
| Sex x ACR x Collection<br>Site | 1 | 1.35 | 1.35 | 0.02 | 0.89 |
| Residuals | 83 | 6137.96 | 73.95 |  |  |
| **ACR vs ALR** |  |  |  |  |  |
| Gen | 1 | 270.14 | 270.14 | 3.39 | 0.07 |
| Cage | 5 | 1496.63 | 299.33 | 3.76 | < 0.005 |
| Sex | 1 | 13842.58 | 13842.58 | 173.93 | < 0.00001 |
| ACR | 1 | 1.87 | 1.87 | 0.02 | 0.88 |
| Sex x ACR | 1 | 154.81 | 154.81 | 1.95 | 0.17 |
| Sex x Collection Site | 1 | 40.27 | 40.27 | 0.51 | 0.48 |
| ACR x Collection Site | 1 | 5.33 | 5.33 | 0.07 | 0.80 |
| Sex x ACR x Collection<br>Site | 1 | 0.32 | 0.32 | 0.00 | 0.95 |
| Residuals | 83 | 6605.77 | 79.59 |  |  |

**Table S6.** ANCOVA of the effect of selection on ASR.

| Effect | DF | SS | MS | F Value | P Value |
| --- | --- | --- | --- | --- | --- |
| **Adult Starvation Resistance (ASR)** |  |  |  |  |  |
| Sex | 1 | 534387.79 | 534387.79 | 3921.59 | < 0.00001 |
| Selection | 1 | 37911.90 | 37911.90 | 278.22 | < 0.00001 |
| Collection Site | 1 | 24.92 | 24.92 | 0.18 | 0.67 |
| Cage | 9 | 9479.13 | 1053.24 | 7.73 | < 0.00001 |
| Generation | 1 | 37157.83 | 37157.83 | 272.68 | < 0.00001 |
| Sex x Selection | 1 | 3087.60 | 3087.60 | 22.66 | < 0.00001 |
| Sex x Collection Site | 1 | 3.82 | 3.82 | 0.03 | 0.87 |
| Sex x Cage | 9 | 387.67 | 43.07 | 0.32 | 0.97 |
| Selection x Generation | 1 | 28129.94 | 28129.94 | 206.43 | < 0.00001 |
| Collection Site x Generation | 1 | 3542.67 | 3542.67 | 26.00 | < 0.00001 |
| Sex x Generation | 1 | 2886.53 | 2886.53 | 21.18 | < 0.00001 |
| Selection x Collection Site x Generation | 1 | 302.40 | 302.40 | 2.22 | 0.14 |
| Sex x Selection x Generation | 1 | 362.82 | 362.82 | 2.66 | 0.10 |
| Sex x Collection Site x Generation | 1 | 0.17 | 0.17 | 0.00 | 0.97 |
| Sex x Selection x Collection Site x Generation | 1 | 527.17 | 527.17 | 3.87 | < 0.05 |
| Residuals | 1047 | 142672.72 | 136.27 |  |  |

**Table S7.** ANCOVA testing the correlation between ACR and ASR.

| Effect | DF | SS | MS | F Value | P Value |
| --- | --- | --- | --- | --- | --- |
| **ACR vs. ASR** |  |  |  |  |  |
| Generation | 1 | 9937.56 | 9937.56 | 115.79 | < 0.00001 |
| Cage | 5 | 1381.00 | 276.20 | 3.22 | < 0.02 |
| Sex | 1 | 55806.79 | 55806.79 | 650.24 | < 0.0001 |
| ACR | 1 | 851.66 | 851.66 | 9.92 | < 0.003 |
| Sex x ACR | 1 | 1.85 | 1.85 | 0.02 | 0.88 |
| Sex x Collection Site | 1 | 4.87 | 4.87 | 0.06 | 0.81 |
| ACR x Collection Site | 1 | 25.28 | 25.28 | 0.29 | 0.59 |
| Sex x ACR x Collection Site | 1 | 86.90 | 86.90 | 1.01 | 0.32 |
| Residuals | 83 | 7123.44 | 85.82 |  |  |

**Table S8.** ANCOVA of the effect of selection on aversion to copper-contaminated food.

| Effect | DF | SS | MS | F Value | P Value |
| --- | --- | --- | --- | --- | --- |
| **Copper Aversion (CA)** |  |  |  |  |  |
| Treatment | 2 | 451.28 | 225.64 | 1599.98 | < 0.00001 |
| Sex | 1 | 5.68 | 5.68 | 40.31 | < 0.00001 |
| Collection Site | 1 | 0.85 | 0.85 | 6.06 | < 0.02 |
| Selection | 1 | 1.15 | 1.15 | 8.16 | < 0.005 |
| Cage | 9 | 5.64 | 0.63 | 4.45 | < 0.00001 |
| Generation | 1 | 43.45 | 43.45 | 308.13 | < 0.00001 |
| Treatment x Sex | 2 | 12.42 | 6.21 | 44.02 | < 0.00001 |
| Treatment x Collection Site | 2 | 0.16 | 0.08 | 0.56 | 0.57 |
| Sex x Collection Site | 1 | 0.07 | 0.07 | 0.52 | 0.47 |
| Treatment x Selection | 2 | 0.70 | 0.35 | 2.49 | 0.08 |
| Sex x Selection | 1 | 0.54 | 0.54 | 3.80 | 0.05 |
| Treatment x Cage | 18 | 2.52 | 0.14 | 0.99 | 0.46 |
| Sex x Cage | 9 | 4.98 | 0.55 | 3.92 | < 0.00006 |
| Treatment x Generation | 2 | 0.77 | 0.39 | 2.74 | 0.06 |
| Collection Site x Generation | 1 | 0.04 | 0.04 | 0.31 | 0.58 |
| Selection x Generation | 1 | 1.02 | 1.02 | 7.20 | < 0.008 |
| Sex x Generation | 1 | 0.00 | 0.00 | 0.01 | 0.92 |
| Treatment x Sex x Collection Site | 2 | 0.47 | 0.23 | 1.66 | 0.19 |
| Treatment x Sex x Selection | 2 | 0.26 | 0.13 | 0.93 | 0.40 |
| Treatment x Sex x Cage | 18 | 1.14 | 0.06 | 0.45 | 0.98 |
| Treatment x Collection Site x Generation | 2 | 0.01 | 0.01 | 0.04 | 0.96 |
| Treatment x Selection x Generation | 2 | 0.02 | 0.01 | 0.09 | 0.92 |
| Collection Site x Selection x Generation | 1 | 0.00 | 0.00 | 0.02 | 0.88 |
| Treatment x Sex x Generation | 2 | 0.39 | 0.20 | 1.39 | 0.25 |
| Sex x Collection Site x Generation | 1 | 0.69 | 0.69 | 4.86 | < 0.03 |
| Sex x Selection x Generation | 1 | 0.14 | 0.14 | 0.97 | 0.33 |
| Treatment x Collection Site x Selection x Generation | 2 | 0.59 | 0.30 | 2.11 | 0.12 |
| Treatment x Sex x Collection Site x Generation | 2 | 0.06 | 0.03 | 0.23 | 0.79 |
| Treatment x Sex x Selection x Generation | 2 | 0.08 | 0.04 | 0.28 | 0.75 |
| Sex x Collection Site x Selection x Generation | 1 | 0.01 | 0.01 | 0.06 | 0.81 |
| Treatment x Sex x Collection Site x Selection x Generation | 2 | 0.08 | 0.04 | 0.27 | 0.76 |
| Residuals | 18536 | 2614.10 | 0.14 |  |  |

**Table S9.** ANCOVA testing the correlation between ACR and aversion to copper in food.

| Effect | DF | SS | MS | F Value | P Value |
| --- | --- | --- | --- | --- | --- |
| **Aversion to 0.5mM (CA0.5)** |  |  |  |  |  |
| Generation | 1 | 0.18 | 0.18 | 8.52 | < 0.005 |
| Cage | 5 | 0.18 | 0.04 | 1.73 | 0.14 |
| Sex | 1 | 0.00 | 0.00 | 0.01 | 0.92 |
| ACR | 1 | 0.05 | 0.05 | 2.60 | 0.11 |
| Sex x ACR | 1 | 0.01 | 0.01 | 0.51 | 0.48 |
| Sex x Collection Site | 1 | 0.04 | 0.04 | 1.81 | 0.18 |
| ACR x Collection Site | 1 | 0.01 | 0.01 | 0.36 | 0.55 |
| Sex x ACR x Collection Site | 1 | 0.01 | 0.01 | 0.73 | 0.40 |
| Residuals | 83 | 1.71 | 0.02 |  |  |
| **Aversion to 1mM (CA1)** |  |  |  |  |  |
| Generation | 1 | 0.16 | 0.16 | 5.21 | < 0.03 |
| Cage | 5 | 0.13 | 0.03 | 0.84 | 0.52 |
| Sex | 1 | 0.07 | 0.07 | 2.27 | 0.14 |
| ACR | 1 | 0.07 | 0.07 | 2.32 | 0.13 |
| Sex x ACR | 1 | 0.01 | 0.01 | 0.31 | 0.58 |
| Sex x Collection Site | 1 | 0.01 | 0.01 | 0.24 | 0.62 |
| ACR x Collection Site | 1 | 0.00 | 0.00 | 0.00 | 0.97 |
| Sex x ACR x Collection Site | 1 | 0.00 | 0.00 | 0.16 | 0.69 |
| Residuals | 82 | 2.52 | 0.03 |  |  |
| **Aversion to 2mM (CA2)** |  |  |  |  |  |
| Generation | 1 | 0.38 | 0.38 | 8.93 | < 0.004 |
| Cage | 5 | 0.30 | 0.06 | 1.42 | 0.23 |
| Sex | 1 | 0.12 | 0.12 | 2.91 | 0.09 |
| ACR | 1 | 0.01 | 0.01 | 0.34 | 0.56 |
| Sex x ACR | 1 | 0.01 | 0.01 | 0.30 | 0.59 |
| Sex x Collection Site | 1 | 0.00 | 0.00 | 0.01 | 0.93 |
| ACR x Collection Site | 1 | 0.08 | 0.08 | 2.00 | 0.16 |
| Sex x ACR x Collection Site | 1 | 0.04 | 0.04 | 1.02 | 0.32 |
| Residuals | 83 | 3.51 | 0.04 |  |  |

**Table S10.** ANCOVA of the effect of selection on average lifespan (ALS)

| Effect | DF | SS | MS | F Value | P Value |
| --- | --- | --- | --- | --- | --- |
| **Average Lifespan (ALS)** |  |  |  |  |  |
| Sex | 1 | 5837.52 | 5837.52 | 106.32 | < 0.00001 |
| Selection | 1 | 5780.81 | 5780.81 | 105.29 | < 0.00001 |
| Collection Site | 1 | 6601.16 | 6601.16 | 120.23 | < 0.00001 |
| Cage | 9 | 10529.55 | 1169.95 | 21.31 | < 0.00001 |
| Generation | 1 | 364.30 | 364.30 | 6.64 | < 0.02 |
| Sex x Selection | 1 | 148.88 | 148.88 | 2.71 | 0.10 |
| Sex x Collection Site | 1 | 15.27 | 15.27 | 0.28 | 0.60 |
| Sex x Cage | 9 | 549.34 | 61.04 | 1.11 | 0.35 |
| Collection Site x Generation | 1 | 111.61 | 111.61 | 2.03 | 0.15 |
| Selection x Generation | 1 | 1221.74 | 1221.74 | 22.25 | < 0.00001 |
| Sex x Generation | 1 | 1114.39 | 1114.39 | 20.30 | < 0.00001 |
| Selection x Collection Site x Generation | 1 | 764.94 | 764.94 | 13.93 | < 0.0003 |
| Sex x Collection Site x Generation | 1 | 109.32 | 109.32 | 1.99 | 0.16 |
| Sex x Selection x Generation | 1 | 128.17 | 128.17 | 2.33 | 0.13 |
| Sex x Selection x Collection Site x Generation | 1 | 1.20 | 1.20 | 0.02 | 0.88 |
| Residuals | 592 | 32503.69 | 54.90 |  |  |

**Table S11.** ANCOVA testing the correlation between ACR and ALS.

| Effect | DF | SS | MS | F Value | P Value |
| --- | --- | --- | --- | --- | --- |
| **ACR vs ALS** |  |  |  |  |  |
| Generation | 1 | 15.24 | 15.24 | 0.70 | 0.41 |
| Cage | 5 | 175.27 | 35.05 | 1.60 | 0.18 |
| Sex | 1 | 803.82 | 803.82 | 36.68 | < 0.00001 |
| ACR | 1 | 263.19 | 263.19 | 12.01 | < 0.002 |
| Sex x ACR | 1 | 37.70 | 37.70 | 1.72 | 0.20 |
| Sex x Collection Site | 1 | 0.52 | 0.52 | 0.02 | 0.88 |
| ACR x Collection Site | 1 | 122.75 | 122.75 | 5.60 | < 0.03 |
| Sex x ACR x Collection Site | 1 | 22.94 | 22.94 | 1.05 | 0.31 |
| Residuals | 47 | 1030.09 | 21.92 |  |  |

## Notes

### Competing Interest Statement

The authors have declared no competing interest.

